# A novel non-catalytic function of PA2803-encoded PcrP contributes to polymyxin B resistance in *P. aeruginosa* and redefines the functional role of the PA2803 subfamily

**DOI:** 10.1101/2025.05.13.653872

**Authors:** T. Salpadoru, S. Khanam, V.A. Borin, Ma. A. Achour, Denise Oh, M. Kanik, P.C. Gallage, A. Khanov, M. Hull, S. Pitre, P.K. Agarwal, M.J. Franklin, M.A. Patrauchan

## Abstract

The opportunistic human pathogen *Pseudomonas aeruginosa* (*Pa*), a leading cause of severe infections, becomes increasingly resistant to antibiotics, including the last resort antibiotic, polymyxin-B (PMB). Previous studies have shown that calcium (Ca^2+^) at the levels encountered during infections increases *Pa* resistance to PMB. However, the mechanisms of this Ca^2+^ regulation are not known. Here, we identified three novel genes (*PA2803, PA3237* and *PA5317)* that contribute to the Ca^2+^-dependent PMB resistance in *Pa. PA2803,* the focus of this work, encodes a putative phosphonatase and is a founding member of the PA2803 subfamily from the Haloacid Dehalogenase Superfamily. Since the transcription of this gene is regulated by both Ca^2+^ and inorganic phosphate (P_i_), we named it “<u>P</u>_i_ and <u>C</u>a^2+^ <u>r</u>egulated <u>p</u>rotein, PcrP”. Congruent with sequence-based predictions, we showed that PcrP lacks catalytic activity and instead binds protein partners, revealing a novel non-catalytic function for PA2803 subfamily proteins. By using pull-down assays and bacterial two-hybrid system, we identified and validated two protein partners of PcrP: Acp3 and PA3518. We show that PcrP is involved in oxidative stress responses in *Pa*, which are likely mediated by its interactions with Acp3, and may support its role in PMB resistance. In addition, PcrP imparts a Ca^2+^-dependent growth advantage to *Pa* during P_i_ starvation and plays a role in polyphosphate accumulation in a Ca^2+^-dependent manner. Overall, this study identified a novel protein-binding function for the PA2803 subfamily representative that mediates *Pa* responses to elevated Ca^2+^ and P_i_ starvation and enhances PMB resistance.

**IMPORTANCE:** *Pseudomonas aeruginosa* (*Pa*) is a critical human pathogen that presents significant clinical challenges, underscoring the urgent need for understanding its resistance mechanisms. Previous studies have shown that calcium (Ca^2+^) at the levels detected during infections increase *Pa* resistance to the last resort antibiotic polymyxin-B (PMB). For the first time, we identified three novel genes, whose products are required for the Ca^2+^-dependent PMB resistance in *Pa.* One of them, *PA2803,* regulated by Ca^2+^ and phosphate, was named <u>p</u>hosphate and <u>C</u>a^2+^ <u>r</u>egulated <u>p</u>rotein, PcrP. This study discovered a novel protein-binding function of PcrP and identified two protein partners. The protein-binding function is likely shared by the entire PA2803 subfamily of proteins, which redefines their previous functional assignment as enzymes.

## INTRODUCTION

To survive, bacteria perceive changes in their environments by using complex regulatory networks that elicit adaptive responses. Some environmental cues can directly or indirectly modulate bacterial antimicrobial susceptibility (1,2). *Pseudomonas aeruginosa* (*Pa*) is a versatile human pathogen known for its ability to adapt to diverse environments, including the human body (3–6). *Pa* is a leading cause of morbidity and mortality in immuno-compromised patients and in patients suffering from cystic fibrosis (CF). It is also well known for its high resistance to nearly all antibiotics, and therefore, regarded as one of the leading pathogens that present a progressively grave risk for global public health (7–10).

The success of *Pa* as a pathogen in a hostile host environment, such as CF lung, relies on successful recognition of host cues and efficient pathoadaptation, which is favored by its high regulatory and metabolic versatility (6,11–13). Two examples of host factors, that also serve as cellular messengers, include calcium (Ca^2+^) and inorganic phosphate (P_i_) (11,14). The levels of these ions within the host can be recognized by invading pathogens and may guide their host adaptation. Our earlier work showed that the elevated levels of Ca^2+^ commonly detected in body fluids of CF patients (11,15,16) trigger alterations in the transcriptome and proteome of *Pa,* including the production of multiple virulence factors, such as pyocyanin, pyoverdine, swarming motility, and biofilm formation, known to enhance the ability of *Pa* to infect its host (11,17–22). Elevated Ca^2+^ has also been shown to induce the acute-to-chronic virulence switch during *Pa* infection (23). Similarly, depleted levels of P_i_ induce the production of pyocyanin and pyoverdine in *Pa* (24–29). Importantly, in response to elevated Ca^2+^ and low P_i,_ *Pa* susceptibility to several antibiotics is increased. These include polymyxin B (PMB), recognized as a “last hope” antibiotic for multi-drug resistant *Pa* infections (14,17,30,31).

Several studies characterized multiple mechanisms of *Pa* resistance to PMB, with most of them involved in modifying the primary target of PMB, lipopolysaccharide (LPS) molecules in the outer membrane (32). Currently, at least six two-component systems (TCSs) are known to regulate PMB resistance: PhoP/Q(33,34), PmrA/B(33,35), ParR/S(36), ColR/S(37), CprR/S(38) and CbrA/B(39). This regulation is primarily *via* controlling the expression of the *arnBCADTEF* operon responsible for introducing phosphoethanolamine (PEtN) and 4-amino-4-deoxy-L-arabinose (L-Ara4N) groups to the lipid A moiety of LPS (32,40,41). However, Ca^2+-^ dependent mechanisms of PMB resistance have not been identified. Here, we present three novel Ca^2+^-dependent players in *Pa* resistance to PMB, including *PA2803.* The gene encodes a putative phosphonoacetaldehyde hydrolase (phosphonatase) and is recognized as the founder of the PA2803 subfamily belonging to haloacid dehalogenase superfamily (HADSF).

HADSF is one of the largest protein superfamilies present in all domains of life (Fig. S1) and playing diverse biological functions (42). Sequence diversity within the family enables diverse catalytic as well as non-catalytic functions (43–45). The majority of the members are enzymes involved in phosphoryl transfer reactions, such as phosphatases, P-type ATPases, phosphonatases, and phosphotransferases (42,46). Despite functional differences, all HADSF members share a conserved catalytic “core” domain featuring four conserved sequence motifs and often contain a “cap” domain that defines substrate specificity and aids in catalysis (42,43,47,48). The presence, position and the topology of this cap domain classify HADSF proteins into three structural classes, C1, C2 and C0 (reviewed in (42)). PA2803, a member of the C1 class, shares similarities with known phosphonatases, but has a truncated cap domain, placing into a unique subfamily exclusive to *Pseudomonads*, termed the “PA2803 subfamily” (49). This subfamily lacks conservation of key active-site residues necessary for magnesium cofactor interaction, substrate binding, and catalysis, indicating a loss of catalytic function (42,49). This was supported by the lack of catalytic activity in one member of the PA2803 subfamily, PSPTO_2114 from *P. syringae pv. tomato* (49). However, no alternative function of the subfamily has been elucidated.

Here we present the first experimental evidence that PA2803 functions by binding protein partners. Given the high level of sequence conservation observed within the PA2803 (PcrP) subfamily (49), the protein-binding function of PcrP is likely shared by other members of the subfamily. Considering PA2803 transcriptional regulation by phosphate and Ca^2+^, we named this protein PcrP (<u>P</u>hosphate and <u>C</u>a^2+^-<u>r</u>egulated <u>p</u>rotein). We propose that PcrP role in Ca^2+^-induced PMB resistance can be mediated by its contribution to oxidative stress response.

## MATERIALS AND METHODS

### Bacterial strains, plasmids and media

Strains and plasmids used in this study are listed in Table 1. *Pa* strain PAO1 is the non-mucoid strain with genome sequence available (50). All the strains were maintained in 10% skim milk at −80°C. For each experiment, bacteria were inoculated from frozen stocks onto LB agar containing the appropriate antibiotic when applicable and grown overnight at 37°C, from which isolated colonies were used to inoculate precultures. *Pa* strains used for pull-down assays were grown in media containing 300 µg/ml carbenicillin (Cb). For bacterial two-hybrid (BTH) assays, *Escherichia coli* BTH101 strains carrying constructs, were grown in Luria-Bertani (LB) (Goldbio) agar supplemented with 40 µg/ml streptomycin (Strep), 50 µg/ml kanamycin (Kan) or 100 µg/ml ampicillin (Amp). 25⁰ C was considered as the room temperature (RT).

**Table 1.**
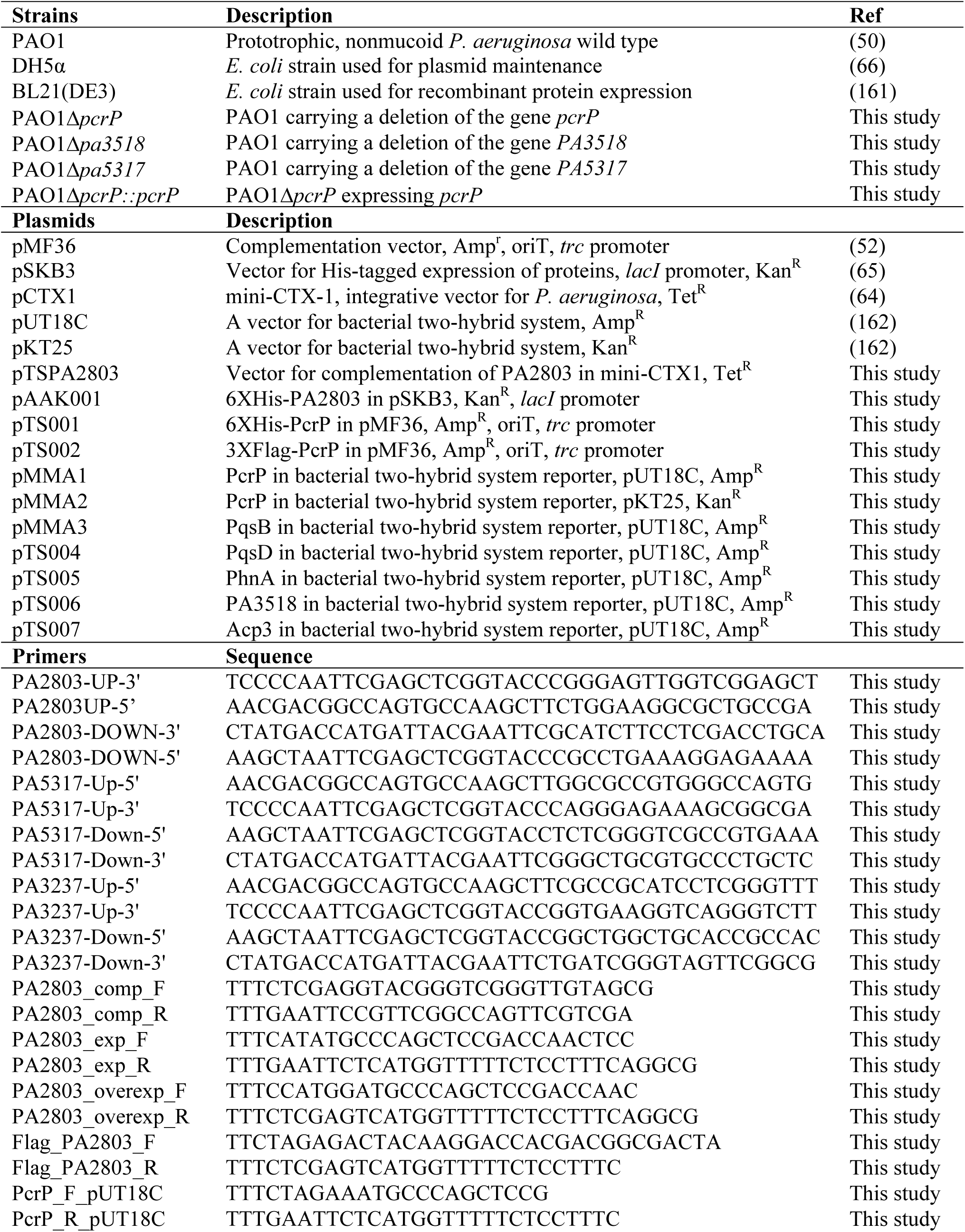
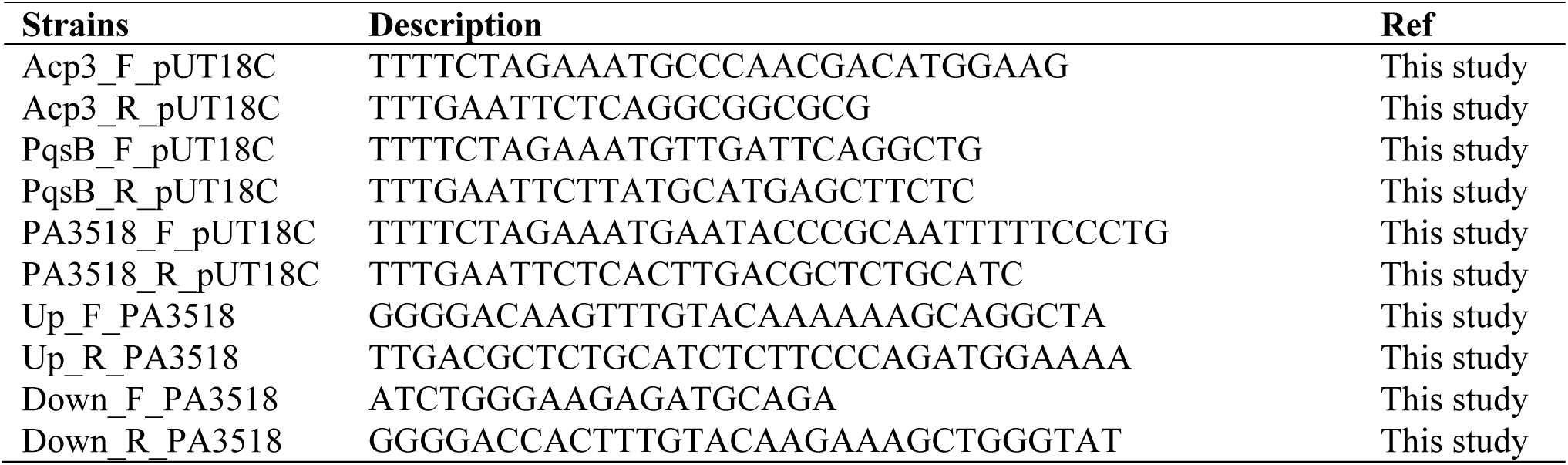
Strains and plasmids used in this study.

For antimicrobial susceptibility and growth analyses at high P_i_, biofilm minimal medium (BMM) (20) was used. BMM contained (per liter) 9.0 mM sodium glutamate, 50 mM glycerol, 0.02 mM MgSO_4_, 0.15 mM NaH_2_PO_4_, 0.34 mM K_2_HPO_4_, and 145 mM NaCl, 200 µl trace metals, 1 ml vitamin solution. Trace metal solution (per liter of 0.83 M HCl): 5.0 g CuSO4.5H_2_O, 5.0 g ZnSO4.7H2O, 5.0 g FeSO4.7H2O, 2.0 g MnCl_2_.4H2O). Vitamins solution (per liter): 0.5 g thiamine, 1 mg biotin. The pH of the medium was adjusted to 7.0. BMM8 medium used for phenotypic assays had the same composition as BMM but contained 0.08 mM MgSO_4_. Low P_i_ medium BMMH8 used for P_i_-starvation experiments contained 9.0 mM sodium glutamate, 50 mM glycerol, 0.08 mM MgSO4, 10 mM HEPES, and 145 mM NaCl, 200 µl trace metals, 1 ml vitamin solution. To vary P_i_ levels, BMMH8 was supplemented with K_2_HPO_4_ at 50 µM (low) and 580 µM (high) P_i_ levels. The media were supplemented with 1.5% agar (Difco) for agar-plate based assays.

### Transposon mutants

Transposon insertion mutants were obtained from the University of Washington Two - Allele library (NIH grant # P30 DK089507) (Table 1). The mutants contained ISphoA/hah or ISlacZ/hah insertions with tetracycline resistance cassette that disrupted the genes of interest. The mutations were confirmed by two-step PCR: first, transposon flanking primers were used to verify that the target gene is disrupted, and second, transposon-specific primers were used to confirm the transposon insertion. The primer sequence is available at www.gs.washington.edu. For convenience, the mutants were designated as PA::Tn5, where PA is the identifying number of the disrupted gene from *Pa* PAO1 genome (www.pseudomonas.com)

### Random mutagenesis, selection of PMB sensitive mutants, and identification of the mutated genes

To identify genes responsible for PMB resistance at elevated Ca^2+^, the *Pa* PAO1 genome was subjected to random mutagenesis with N-methyl N-nitro-N-nitrosoguanidine (NTG) using the method described in (51). First, *Pa* was plated on BMM with no added Ca^2+^ or with 5 mM CaCl_2_, supplemented with PMB at 2 to 64 μg/ml to identify PMB concentrations that were permissive for growth in the two media. From these experiments, we identified 8 μg/ml PMB as a cutoff concentration that was permissive for *Pa* growth in 5 mM CaCl_2_, but not in BMM with no added CaCl_2_. Next, *Pa* was mutagenized with NTG by exposing log-phase cultures grown in LB medium to 50 μg/ml NTG for 30 min with incubation at 37^0^C. Cells were washed with saline, then serially diluted in saline to achieve a cell concentration that gave approximately 100-200 colonies per plate. Approximately 8000 colonies were plated on BMM + 5 mM CaCl_2_ with no antibiotic. The colonies were replica plated onto BMM supplemented with 5 mM CaCl_2_ and 8 μg/ml PMB. About twenty colonies that grew on BMM + 5 mM CaCl_2_ but did not grow on BMM + 5 mM CaCl_2_ with 8 μg/ml PMB were selected for further analysis. The isolated cultures were streaked on BMM + 5 mM CaCl_2_ and 8 μg/ml PMB to ensure lack of growth. Three colonies that had the lowest MICs of PMB in 5 mM CaCl_2_ were selected for further study.

Genes responsible for PMB resistance at 5 mM CaCl_2_ were identified by complementing the mutant strains with a randomly fragmented *Pa* PAO1 gene bank. For gene bank construction, genomic DNA from *Pa* PAO1 was partially digested with FatI (CATG recognition sequence) (NEB) then ligated into the NcoI restriction site (with compatible cohesive ends) of the *Pa* expression vector, pMF36 (52). *E. coli* HB101 was transformed with pMF36/random gene bank generating approximately 10,000 clones. The colonies were pooled, and the pooled colonies were conjugated into the *Pa* PMB mutant strains using the conjugation helper plasmid pRK2013. The conjugation mix was plated on BMM containing 5 mM CaCl_2_, 150 μg/ml of Cb to select for the pMF36 plasmid, and 8 μg/ml PMB. Plasmid DNA was isolated from the colonies with restored resistance to 8 μg/ml PMB. Genes were identified by plasmid sequencing.

### Antibiotic susceptibility assays

*Pa* resistance to PMB was assayed as described in (53). Briefly, bacterial strains were grown in BMM at no added or supplemented with 5 or 10 mM CaCl_2_. 100 µl of the mid-log cultures (12 h as defined by OD_600_-monitored growth curves) normalized to the OD_600_ of 0.1 were spread inoculated onto the surface of BMM agar without or supplemented with the appropriate CaCl_2_. E-test strips for PMB (Biomeurix) were placed on the surface of the inoculated plates and incubated for 24 h. The minimum inhibitory concentration (MIC) was recorded according to the manufacturer’s instructions.

### RNA sequencing (RNA-seq)

*Pa* PAO1 was grown to mid-log growth phase (12 h) in BMM with or without 5 mM CaCl_2_. Bacterial cells were collected, and RNA was isolated as previously described (54). Briefly, total RNA, in three replicates for each growth condition, was isolated by using RNeasy Protect Bacteria Mini kit (Qiagen) or ZR Fungal/ Bacterial RNA MiniPrep TM (Zymo Research). The quality of the purified RNA was assessed by Bioanalyzer 2100 (Agilent) and 1% agarose gel electrophoresis. RNA-Seq analysis was performed at Vertis Biotechnology AG (Germany) (55). Prior to library preparation, ribosomal RNA was removed from the total RNA samples using Ribo-Zero rRNA Removal Kit for bacteria (Illumina). The RNA library was amplified and sequenced on an Illumina NextSeq 500 system using 75 bp read length. The NCBI accession number is PRJNA703087. The Illumina reads were aligned to the reference *Pa PAO1* genome (downloaded from the Pseudomonas Genome Database) with 95-97% alignment rate by using the short read alignment program Bowtie 2 (56). Quantification of transcripts was achieved by using RSEM (RNA-seq by Expectation-Maximization) program (57). Statistical analysis for differential gene expression was conducted using the RSEM count matrixes in EdgeR (58). Differently expressed genes were selected based on a log_2_ fold change (log_2_FC) ≥ |1.0| and false discovery rate (adjusted p value ≤ 0.05). For additional gene annotation, homology searches using NCBI and UniProt databases were used.

### Sequence analysis and protein structure predictions

To infer phylogenetic relationships of PA2803 (below designated as PcrP), its amino acid sequence was aligned with a curated list of homologs identified by a BlastP search using UniProt and PDB databases. The alignment was done using MUSCLE with default alignment parameters, and the evolutionary history was inferred by using the Maximum Likelihood and JTT matrix-based model [2] in MEGA11 [3] with the bootstrap consensus tree inferred from 50 replicates [4]. Branches reproduced in less than 50% bootstrap replicates were collapsed. Initial tree(s) for the heuristic search were obtained automatically by applying Neighbor-Join and BioNJ algorithms to a matrix of pairwise distances estimated using the JTT model and then selecting the topology with superior log likelihood value.

### Construction of deletion mutations and genetic complements

Deletion mutants were generated by allelic exchange using Seamless Ligation cloning (SLICE) (59–61). Approximately 1kb of the upstream DNA of the *PA2803* gene was PCR amplified using Phusion High fidelity polymerase (New England Biolabs #M0530S) and primers: PA2803-UP-3’, PA2803UP-5’, and approximately 1 kb of downstream DNA was amplified with primers: PA2803-DOWN-3’ and PA2803-DOWN-5’ (Table 1). The PCR products and the gentamicin (Gm) resistance gene from pFGM1 (59,62) were assembled into the gene replacement vector, pEX18 (60,63). SLICE reaction (10 μl) contained 100 ng of pEX18 digested with EcoRI and HindIII, the Gm resistance fragment digested with KpnI, and the two PCR products diluted to be 1:3 molar ratio of the pEX18 fragment. The SLICE reaction was carried out at 37 °C for 60 min, then 2.5 μl of the reaction was transformed into *E. coli* TOP10 cells. Gm-resistant colonies with the correctly assembled gene replacement plasmid were confirmed by colony PCR, restriction digest, and Sanger sequencing at the OSU Core Facility. The plasmid was then electroporated into *P. aeruginosa* PAO1 and colonies selected on Gm^50^ plates for merodiploids. Colonies were then cultured in 3ml LB medium containing 50 ug/ml Gm and streaked on Gm^50^ plates with 10% sucrose to obtain allelic exchange mutants. The mutant strains were tested for Cb sensitivity and by PCR to verify allelic exchange. Deletion mutants of genes *PA3237* and *PA5317* were generated using the same approach using the respective primer pairs PA3237-UP-3’, PA3237-UP-5’; PA3237-DOWN-3’, PA3237-DOWN-5’, and PA5317-UP-3’, PA5317-UP-5’; PA5317-DOWN-3’, PA5317-DOWN-5’ (Table 1).

The Δ*pcrP* mutant was complemented as described in (64) with few modifications. Using Q5 High-Fidelity DNA polymerase (NEB) and the primer pair PA2803_compF and PA2803_compR (Table 1), the PAO1 genomic region that includes *pcrP* (744 bp) and the 375-bp upstream and 40-bp regions downstream of the gene were amplified. The amplicon was digested with the restriction enzymes, XhoI and EcoRI, and ligated into mini-CTX1; and the insertion was confirmed by Sanger sequencing at the OSU Core Facility. The resulting plasmid, pTSPA2803 was transformed into *E. coli* SM10 by heat shock. The miniCTX construct, pTSPA2803, was incorporated into the *P. aeruginosa* Δ*pcrP* genome by conjugation. The transconjugants were selected on PIA agar plates containing 60 µg/ml of Tet and verified by PCR using the PA2803_compF/ PA2803-compR primers.

### Cloning, expression, and purification of PcrP

*Pa* PAO1 genomic DNA was extracted using the Wizard Genomic DNA Purifcation Kit (Promega) and used as the template to PCR amplify the 744 bp DNA fragment containing *pcrP* (PA2803) using Q5 high-fidelity DNA polymerase (NEB Inc.) and primers PA2803_exp_F and PA2803_exp_R (Table 1) with restriction sites NdeI and EcoRI, respectively. The resulting PCR product was column-purified (Zymo) and cloned into pSKB3 vector (65) using Quick Ligase (NEB Inc.). The ligation product was transformed into *E. coli* DH5α (66), and Kan-resistant transformants were selected and verified by Sanger sequencing. The verified construct pAAK001 carrying 6xHis-PcrP (Table 1) was heat-shock transformed into *E. coli* BL21 (DE3) cells (Novogene). BL21 cells carrying 6XHis-PcrP were grown in 5 ml LB containing 50 µg/ml Kan at 37 ⁰C with agitation at 200 rpm and used to inoculate fresh 1 L of LB-Kan at a ratio of 1:100. The cultures were grown to an OD_600_ of 0.6–1.0 and induced by adding 15 µM (final) isopropyl a-D-thiogalactopyranoside (IPTG) and incubating for additional 16 h at 30 ⁰C. The cells were harvested by centrifugation at 5,514 g for 15 min at 4 ⁰C, and the resulting cell pellet was resuspended in 50 ml of ice-cold buffer A (20 mM Tris, 500 mM NaCl, 20 mM imidazole, 1% glycerol, pH 7.8 at 25⁰C). Lysozyme was added to the cell suspension at 0.5 mg/ml, incubated for 40 min at 4 ⁰C, and the cells were sonicated for a 100 s in cycles of 20 s/on and 20 s/off using a Fisher ultrasonic processor XL 2010 at power setting of 5. The lysate was collected by centrifugation at 9,803 g for 30 min at 4 °C. The supernatant was loaded onto a gravity-flow column containing 4 ml of HisPur Nickel Nitrilotriacetic acid resin (Ni–NTA) (Thermofisher) pre-equilibrated with buffer A. The column was rocked for 1 h at 4 °C to facilitate binding. The Ni–NTA resin was subsequently washed with 100 ml of ice-cold buffer A, and the protein was eluted with 20 ml buffer B (20 mM Tris, 500 mM NaCl, 1% glycerol, 250 mM Imidazole, pH 7.8). The eluted fractions were analyzed by SDS-PAGE, and those containing the protein with desired MW were combined, dialyzed against buffer A, concentrated at 4 °C using a 10K Amicon Ultra Centrifugal filter (Millipore), and flash frozen in liquid N_2_ to be stored at −80 °C. The protein concentration was determined by Bradford method (67) using Quick Start™ Bradford 1x Dye Reagent (Bio-Rad) based on absorbance at 280 nm (Ԑ, 33710 M^−1^ cm^−1^, determined using ExPasy).

### *In-vitro* crosslinking assays of PcrP

Purified PcrP was subjected to crosslinking assays with increasing concentrations of glutaraldehyde. 20 µl reaction mixtures contained 50 µM PcrP, and glutaraldehyde at 0.02, 0.05, 0.1 and 0.2% concentrations in buffer 3 (50 mM NaCl, 1% glycerol in HEPES 20 mM, pH 7.8). These were incubated for 2 min, after which 2 µl of 1M Tris-HCl buffer (pH 8.0) was added to the mixture to terminate the reaction. The mixtures were then mixed with SDS loading buffer and separated on a 12% SDS-PAGE gel to be visualized by staining with Coomassie brilliant blue.

### Synthesis of the sodium salt of phosphonoacetaldehyde (Na-Pald)

The synthesis was performed in two steps. First, diethyl (2-oxoethyl)-phosphonate was synthesized. For this, diethyl 2,2-diethoxyethylphosphonate (1.27 g, 5 mmol) was dissolved in HCl (2M, 15 ml), and the reaction mixture was stirred at RT for 20 h. The aqueous mixture was extracted with EtOAc (4×10 ml), and the organic layer was dried over Na_2_SO_4_. The solvent was removed by vacuum and the crude aldehyde product was purified by column chromatography to obtain a colorless liquid. (441 mg, 2.5 mmol, 50%). The product was stored at −20 °C. The NMR spectrum of the product is shown in Fig. S2A.

Second, the synthesized diethyl (2-oxoethyl)-phosphonate (360 mg, 2 mmol) was dissolved in 1,2-dichloroethane (DCE, 15 ml) under Argon at RT. Allyltrimethylsilane (457 mg, 4 mmol) and Bromotrimethylsilane (1.84 g, 12 mmol, 1.58 ml) were added successively, and reaction mixture was stirred at RT for 24 h. All volatiles were removed in *vacuo* at RT, and the residue was dissolved in DCE (5 ml). The solvent was removed, and the procedure was repeated twice. The remaining liquid was dissolved in water (5 ml), and the pH of the solution was adjusted with NaOH (1M) to 4.0. The yellow/orange solution was lyophilized to obtain a yellowish powder (containing about 0.25 mmol/ 42 mg of Pald per 100 mg) in quantitative yield. The NMR spectrum of the product is shown in Fig. S2B.

### Enzymatic assays

Phosphonatase catalytic activity of PcrP was measured as previously described (47). Briefly, 6XHis-PcrP was used at a concentration of 50 µM in a standard assay solution containing 1 mM Pald (synthesized as described above), 5 mM MgCl_2_, 0.13 mM NADH (Sigma-Aldrich), and 9 units/ml alcohol dehydrogenase (Sigma-Aldrich) in 50 mM HEPES (pH 7.0, 25 °C, 30 °C, or 37 °C). To determine the level of NADH (ε = 6200 M^−1^ cm^−1^) in the reaction medium absorbance was monitored at 340 nm 30 min with reads every 2 min. A reaction mixture that contained alcohol dehydrogenase, NADH and acetaldehyde (Thermo Fisher Scientific) was used as the positive control.

Phosphatase activity of PcrP was measured by using p-nitrophenyl phosphate (pNPP) as a substrate. The rate of pNPP hydrolysis was determined by monitoring the increase in absorbance at 410 nm (ε = 18.4 mM^−1^ cm^−1^) at 25 °C as previously described (49). Briefly, absorbance at 410 nm was monitored in 1 ml assay mixtures containing 10 µM PcrP, 50 mM HEPES (pH 7.0), 5 mM MgCl_2_, and 1 mM pNPP (Sigma-Aldrich) for 30 min at RT. A reaction mixture containing 1 unit of shrimp alkaline phosphatase (New-England Biolabs) was used as the positive control.

### Identification of PcrP binding partners by pull-down assays

To identify PcrP binding protein partners, Ni-based and co-immunoprecipitation-based pull-down assays were conducted using 6x-His and 3xFlag-PcrP as bait. To generate 6x-His-PcrP construct, the DNA fragment encoding 6x-His tagged *pcrP* obtained by digesting pAAK001 with Xba1 and Xho1 was subcloned into pMF36 (52) to generate pTS001. The plasmid pTS001 was then transformed into *E. coli* DHα by heat shock followed by selection on LB-Amp plates. Transformants were verified by Sanger sequencing. Upon verification, pTS001 plasmid was transformed into Δ*pcrP* by electroporation. For 3xFlag tagging, the DNA fragment containing 3xFlag-*pcrP* was amplified from PAO1 genomic DNA using primers Flag_PA2803_F and Flag_PA2803_R (Table 1) and Q5 DNA polymerase (New England Biolabs). The PCR product was cloned between Xba 1 and Xho1 sites into pMF36 vector to yield pTS002, which was transformed into *E. coli* DHα by heat shock, and successful transformants were selected on LB-Amp plates. Sequence verified pTS002 was then transformed into Δ*pcrP* by electroporation.

To proceed with pull-downs, two Δ*pcrP* strains each overexpressing either 6xHis tagged PcrP or 3xFlag tagged PcrP were inoculated into 5 ml of BMM8 (or BMMH8 for low P_i_ conditions) at 0 mM or 5 mM Ca^2+^, and grown for 12 h at 37°C, shaking at 200 rpm. These precultures were normalized to an OD_600_ of 0.3 and inoculated at 1:1000 ratio into 100 ml BMM8 (or BMMH8) at the corresponding CaCl_2_ level. After a 12 h period of incubation, cells were harvested by centrifuging at 9,820 g for 15 min at RT. The collected cell pellets were washed with 10 mM Tris-HCl buffer (pH 7.5) and resuspended in 1 ml of lysis buffer (20% sucrose, 1x Proteinase Inhibitor Cocktail, 0.5 mg/ml lysozyme in 10 mM Tris-HCl pH 7.5) and incubated for 15 min at 4°C. Then, cells were lysed by sonicating with a Fisher ultrasonic processor XL 2010 at power setting of 5 for 100 seconds (5 x 20s on/off cycles) at 4°C and spun at 15,344 g for 10 min at 4°C to separate the supernatant containing cell lysates. These lysates were subjected to further clarification by centrifugation after which, the total protein content was quantified using Bradford assay (67). Cell lysates containing 1 mg/ml total protein was loaded onto resin containing 20 µl of HisPur™ Ni-NTA Resin (Thermo Fisher Scientific) or ANTI-FLAG® M2 Magnetic Beads (Sigma) pre-equilibrated with wash buffer (10 mM Tris-HCl at pH 7.5). To facilitate protein binding, these mixtures were incubated at 4°C for 2 h with gentle rocking. Then, the resins were washed thoroughly with the wash buffer to remove any unbound proteins. After the final wash, the resins were stored at −20°C. LC-MS/MS analysis was performed at the OSU Proteomics Facility. As a negative control, Δ*pcrP* strain grown at respective CaCl_2_ concentration was used. Three replicates of each sample were subjected to LC-MS/MS analysis.

Following LC-MS/MS, the data analyses included the following steps. After removing reverse and contaminant sequences, the label-free <u>q</u>uantification (LFQ) protein intensities were filtered to only include proteins with q-values below 0.01. To enable calculating fold differences, zero values were imputed by the lowest intensity in the dataset. Then, the reads for each identified protein were averaged between three replicates, normalized by the corresponding averaged values detected in the Δ*pcrP* control, and the ratio presented as log_2_. Proteins predicted to localize in the outer membrane, extracellular space or periplasm were excluded due to their unlikely interactions with cytoplasmic PcrP. To restrict selection and to avoid tag bias, proteins identified by both 3xFlag-PcrP and 6X-PcrP with log_2_(test/Δ*pcrP*) > 5 were selected for further steps. Since higher abundance of PcrP observed at 5 mM CaCl_2_ may affect the abundance of protein partners, proteins with abundance of at least 25% of that of the bait were selected as the top candidates.

### Bacterial two-hybrid (BTH) assays

To validate PcrP interacting partners, BTH assays were conducted using the BACTH System kit (Euromedex). The *Pa* PAO1 genes *pcrP, pqsB*, *pqsD, phnA, PA3518* and *acp3* were cloned into the vector pUT18C, while *pcrP* was cloned into pKT25 to be used as bait. Each combination of pUT18C and pKT25-based constructs carrying *pcrP* and a putative binding partner was co-transformed to *E. coli* BTH101, and transformants were selected by plating on LB-Kan-Amp plates. Colonies co-transformed with the vector controls pUT18C and pKT25 served as a negative control. The plasmids pKT25-zip and pUT18C-zip, each carrying a fusion of the leucine zipper of GCN4 were co-transformed to be used as a positive control as per manufacturer’s instructions (68). Transformants carrying both vectors were then used to inoculate 2 ml of LB supplemented with 0.5 mM IPTG, Kan, and Amp. These cultures were grown overnight at 30°C shaking at 200 rpm. For qualitative detection of protein-protein interactions, blue-white colony screening was used. For this, 2 µl of the cultures from the previous step was spotted on LB plates containing 40 μg/ml 5-bromo-4-chloro-3-indolyl-β-D-galactopyranoside (X-gal) (ThermoFisher Scientific) and 0.5 mM IPTG. These plates were incubated at 30°C for 24-48 h, and colonies were checked for their color. The formation of a blue colony in the presence of X-gal was indicative of positive protein-protein interaction, whereas white color indicated no interaction. As an additional negative control, the combination of the vector carrying *pcrP* (pUT18C: *pcrP*) and the counter-vector control (pKT25) were tested as well. For quantitative evaluation of the qualitatively detected protein interactions, the β-galactosidase activity of the corresponding pairs was quantified. For this, 200 µl of the overnight cultures were subjected to Miller assays in clear 96-well plates as previously described (69). Each combination of interacting partners was tested in triplicate and β-galactosidase activity of each replicate was calculated and converted into Miller units according to published methods (69).

### Computational model generation

Experimental structures available for the apo state of Acp3 (PDB ID: 2LTE) and PA3518 (PDB ID 3BJD) were used for building computational models. The experimental structure of PcrP is not available, therefore, it was predicted by using AlphaFold algorithms (https://alphafold.ebi.ac.uk). Different homo-dimeric, hetero-dimeric, tetrameric (dimer of homo-dimers) and hexameric (dimer of homo-dimers and two separate Acp3 chains) complexes were also generated with AlphaFold (70) using the multimer model (71).

### Molecular dynamics (MD) simulations

MD simulations were performed for a total of 8 complexes (Table 2) in explicit water solvent, using a protocol similar to our previous studies (72,73). Model preparation and simulations were performed using the AMBER v16 program suite for biomolecular simulations (74). AMBER’s *ff14SB* force-fields were used for all simulations (75). MD simulations were performed using NVIDIA graphical processing units (GPUs) and AMBER’s *pmemd.cuda* simulation engine using our lab protocols published previously (76,77). We have verified the suitability of ff14SB for simulations at microsecond timescales for a number of proteins complexes (72,73). Standard parameters from ff14SB force-field were used for protein residues and nucleotides. SPC model was used for water (78,79). The AMBER parameters for the counter-ions were used, as available in the *frcmod.ionsjc_spce* and *frcmod.ionslrcm_hfe_spce* files.

**Table 2.** Summary of MD simulations. Structures of PcrP (Alphafold prediction), PA3518 (PDB ID: 3BJD) and Acp3 (PDB ID: 2LTE) were subjected to MD simulations as described in methods. The combinations of proteins/complexes, no. of replicates and the duration of the stimulation are listed along with the stability determined by the energy analyses.

| Protein(s) | Partner(s) | Duration of MD | Stability |
| --- | --- | --- | --- |
| PcrP | PcrP | 1 μs | Stable |
| PA3518 | PA3518 | 1 μs | Stable |
| Acp3 | Acp3 | 1 μs | Stable |
| PcrP-PcrP | PA3518-PA3518 | 1 μs | Stable |
| PcrP-PcrP | Acp3-Acp3 | 1 μs | Stable |
| PcrP-PcrP | Acp3-Acp3, PA3518-Acp3 | 1 μs | Unstable |
| PcrP-PcrP: Acp3-Acp3 | PA3518-PA3518 | 1 μs | Stable* |

\*AlphaFold conformation unstable, manual complex stable (see text).

Starting with the complex structures obtained from AlphaFold, the missing hydrogen atoms were added with the AMBER’s *tleap* program. After processing the protein and substrate coordinates, all systems were neutralized by addition of counter-ions, and the resulting systems were solvated in a rectangular box of SPC/E water, with a 10 Å minimum distance between the protein and the edge of the periodic box. The prepared systems were equilibrated as described previously (80,81). The equilibrated systems were then used to run 1.0 μs of production MD under constant energy conditions (NVE ensemble). The use of NVE ensemble was preferred as it offers better computational stability and performance (82,83). The production simulations were performed at a temperature of 300 K. As NVE ensemble was used for production runs, these values correspond to the initial temperature at start of simulations. The temperature adjusting thermostat was not used in simulations, and over the course of 1.0 μs simulations, the temperature fluctuated around 300 K with RMS fluctuations between 2-4 K, which is typical for well equilibrated systems. A total of 1,000 conformational snapshots (stored every 1,000 ps) collected for each system was used for analysis. Structural visualization and analysis were performed using UCSF Chimera software (84).

### Energy of interaction

The interaction energies were used as a quantitative measure of stability of protein complexes. A significant change in interaction energies over the course of

MD simulations indicated an unstable complex. The energy for interactions was calculated as a sum of electrostatic and van der Waals energy between atom pairs following the protocol previously developed to investigate protein-protein complexes and protein-substrate systems (81,85).

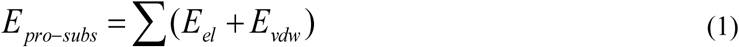

*E_el_* is the electrostatic contribution, *E_vdw_* is the van der Waals term and the summation runs over all atom pairs for the protein-substrate complex. The *E_el_* and *E_vdw_* terms were computed as follows,

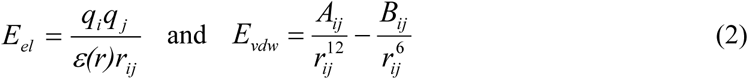

where *q_i_* are par}tial charges, and *A_ij_, B_ij_* are Lennard-Jones parameters. These parameters were obtained from the AMBER force field. A distance-dependent dielectric function was used:

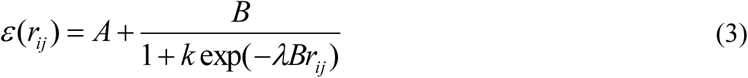

*B*=*ε_o_-A*; *ε_o_*=78.4 for water; *A*= −8.5525; *λ*=0.003627 and *k*=7.7839.

All the atom pairs in complexes were included in the calculations and resulting interaction energies were summed up per residue pair. The energies were calculated for 1,000 snapshots, every 1 ns, sampled during the full 1.0 μs simulation and were averaged over these 1,000 snapshots.

### Measuring endogenous ROS accumulation

To assess ROS accumulation, PAO1 and Δ*pcrP* strains were grown in BMM8 supplemented with or without 5 mM CaCl_2_ for 12 h (mid-log phase). Bacterial cultures (2 ml) were harvested by centrifuging at 2348 g for 5 min. These cells were washed with PBS, resuspended in 1 ml of PBS (pH 7) and normalized to an OD_600_ of 0.5 in two 1.5 ml tubes for DMSO-control and test samples. The ROS indicator, CM-H_2_DFFA (Invitrogen) (1 mM in DMSO), was added to the test samples at a final concentration of 10 µM, while an equal volume of DMSO was added to the control. Bacterial cells were incubated for 30 min at 37 ^0^C in the dark and collected by centrifugation as above. The collected cells were resuspended in 0.5 ml of PBS, and 200 µl of bacterial cell suspensions as well as the PBS control were placed in a black 96-well plate (Corning). OD_600_ and fluorescence at 495/520 nm (Ex/Em) were monitored for 30 min using a Biotek Synergy-Mx plate reader. Each experiment included at least three independent biological replicates per condition. For each replicate, the fluorescence values were normalized by cell density and percent changes in ROS in Δ*pcrP* strain vs PAO1 was calculated for each replicate and then averaged for statistical analysis.

### Phosphate starvation growth studies

To control phosphate levels in the medium, BMM8 was modified into BMMH8 by replacing phosphate with HEPES buffer. BMMH8 contained (per liter): 9.0 mM sodium glutamate, 50 mM glycerol, 145 mM NaCl and 0.08 mM MgSO4 in 10 mM 4-(2-hydroxyethyl)-1-piperazineethanesulfonic acid (HEPES) buffer at pH 7 and was supplemented with 200 µl of trace metals and 1 ml of vitamin solution (described above). When appropriate, K_2_HPO_4_ was added to a final concentration of 50 µM (low P_i_) or 0.58 mM (high P_i_). The high P_i_ level (0.58 mM) equals to that in BMM (26). BMMH8 was supplemented with 5 mM CaCl_2_ when necessary. The precultures (12 h) inoculated from single colonies recovered from frozen stocks were grown in low Pi-BMMH8 to reduce the intracellular Pi storage. These were normalized to OD_600_ of 0.3 and inoculated into 100 ml of fresh BMMH8 medium (1:1000) at low or high P_i_ with 0 or 5 mM CaCl_2_. The cultures were grown shaking at 200 rpm at 37°C for 40 h and monitored for OD_600_.

### Quantification of P_i_ uptake

To assess the role of PcrP in Pi uptake, cultures of PAO1, Δ*pcrP,* and Δ*pcrP*::*pcrP* strains were inoculated and grown as described above, 1 ml samples were collected every 3 h during growth and analyzed for OD_600_ and concentration of P_i_. The latter was quantified by Malachite green assay as described in (26) with minor modifications. Briefly, cell supernatants were collected by centrifuging at 9391 g for 5 min at RT, and 200 µl of these were mixed with the malachite green (BIOMOL® Green, Enzo Life Sciences) as per manufacturer’s instructions and incubated for 30 min at RT. Absorbance at 660 nm was measured by using a Biotek SynergyMx plate reader. BMMH8 medium containing 0-600 mM of K_2_HPO_4_ was used for a standard curve. To calculate P_i_ uptake, the concentration of remaining P_i_ in cell supernatants was subtracted from the P_i_ concentration detected in the fresh medium, and this value was normalized by the corresponding cell density measured at 600 nm.

### Quantification of pyoverdine

Pyoverdine production was assessed by monitoring its florescence (86). Bacterial cultures were grown at low P_i,_ as described above. Samples (200 µl) were collected every 3 h during growth, placed into a black 96-well plate (Corning), and both the fluorescence at 400/460 nm (Ex/Em) and OD_600_ were measured by using Biotek Synergy-Mx. To calculate the abundance of pyoverdine, the obtained fluorescence values were normalized by the OD_600_ of the corresponding cultures.

### Extraction and quantification of pyocyanin

Pyocyanin production was quantified by using chloroform-HCl method as described in (26) with modifications. Briefly, bacterial cultures were grown for 38 h at low P_i_ as described above, and samples (1.5 ml) were collected every 4 h and centrifuged at 15,344 g for 10 min at RT. The obtained supernatants (1 ml) were then transferred into new tubes, and pyocyanin was extracted by vigorous mixing with 1 ml of chloroform. This mixture was allowed to separate, and the bottom solvent phase was collected and extracted by vigorous mixing with 1/3 volume of 0.2 M HCl. The top pink layer containing pyocyanin was separated and quantified by measuring the absorbance at 520 nm in a clear 96-well plate using a Biotek Synergy-Mx plate reader. The values were normalized by the OD_600_ measured prior to centrifugation.

### Extraction of phospholipids and thin-layer chromatography (TLC)

To evaluate changes in the ornithine lipid formation, total lipids were extracted from WT and Δ*pcrP* strains and subjected to TLC separation. For this, bacterial strains were grown in 50 ml of low P_i_ medium (BMMH8) with or without 5 mM Ca^2+^ for 38 h (stationary phase) and harvested by centrifuging at 15,344 g for 10 min at RT. The cell pellets were washed in PBS, adjusted to the same OD_600_, and phospholipids were extracted using the Bligh-Dyer method (87). The final product in chloroform was dried under N_2_ gas and suspended in 100 µL of chloroform.

A silica gel TLC plate (Sigma-Aldrich) was outlined with a baseline 1 inch from the base of the plate and 6 evenly spaced marks. The obtained lipid extracts (10 µl) were spotted using TLC capillary tubes at the baseline on their respective labeled marks. The TLC chamber was set up using 1-propanol: chloroform: ethyl acetate: methanol: water (25: 25: 25: 10: 5.9) solvent system. The spotted TLC plate was developed in the solvent chamber. Immediately after removing the TLC plate from the chamber, the solvent front line was drawn on the plate. Visualization of ornithine lipids was achieved by spraying with the ninhydrin reagent followed by heating for 5-7 min at ∼105⁰ C in the oven. Black-purple-colored spots on the plate indicated the presence of amino lipids in the samples.

### Quantification of polyphosphate (poly P) granule formation

DAPI blue fluorescent staining, typically used for visualizing DNA (361/497 nm Ex/Em), can also be used to visualize and quantify polyP complexes with yellow fluorescence (415/550 nm Ex/Em) The assay was conducted as previously described in (88), using the DAPI-equivalent dye Hoechst 33342, with modifications. To ascertain the validity of the Hoechst-based poly P quantification method, standard curves were generated using both DNA and poly P standards (Thermo, cat # 390932500) and measuring fluorescence of each set of standards at wavelengths specific for DNA and poly P. The poly P standards (ranging from 0.1-100 µg/mL) generated a linear slope with an R^2^ of 0.93 when measured at wavelengths specific for poly P, while the same standards generated a slope with an R^2^ of 0.35 when measured at wavelengths specific for DNA. This indicates that the presence of poly P can be evaluated by using Hoechst without the interference of DNA. Bacterial cultures were first grown in high P_i_ (580 µM) BMM8 at no added Ca^2+^ for 12 h (mid-log) at 37°C shaking at 200 rpm and normalized to OD_600_ of 0.3. These cultures were used to inoculate 50 ml of BMM8 with no or 5 mM CaCl_2_ at 1:1000 ratio. The cultures were grown for 12 h (mid-log) under the same conditions, and cells were subsequently harvested by centrifuging at 2,348 g for 10 min. After one PBS wash, cells were normalized to OD_600_ of 0.5, and 500 µl of thus prepared cell suspension was stained with 100 µl of NucBlue™ Live ReadyProbes™ Reagent (Hoechst 33342) (Invitrogen) for 30 min at RT. This mixture (200 µl) was used to measure fluorescence at 415/ 550 nm (Ex/Em) in a black 96-well plate (Corning) using the Biotek Synergy-Mx plate reader. To calculate the abundance of poly P, the obtained fluorescence values were normalized by cell density measured at 600 nm.

### Statistical analysis and rigor

All the quantitative assays were performed based on three independent biological replicates. Every experiment was repeated at least two-three times for consistency. Significance was determined by one-way and two-way ANOVA (Microsoft Excel v. 16.54 or Prism 10.0) with cutoff of *p*< 0.05.

## RESULTS

### Ca^2+^ increases polymyxin B (PMB) resistance in *Pa*

Elevated Ca^2+^ has been reported by us and other groups to enhance the resistance of *Pa* to several antibiotics, including PMB and colistin (30,89–91). To investigate Ca²⁺-induced resistance to PMB, we first assessed the PMB minimum inhibitory concentration (MIC) in *Pa* strain PAO1 grown in BMM with or without 5 mM Ca²⁺ and tested for PMB susceptibility using E-test strips. The concentration of Ca^2+^ was chosen to represent the ion levels commonly detected in saliva and nasal secretions in CF patients (e.g. 4.8 ± 0.7 mM in saliva of CF patients *vs* 0.3 mM Ca^2+^ in healthy individuals (15)). Consistent with previous studies, we show that the presence of 5 mM Ca^2+^ enhances *Pa* resistance to PMB by 12-fold (Fig. 1A).

**Figure 1.**
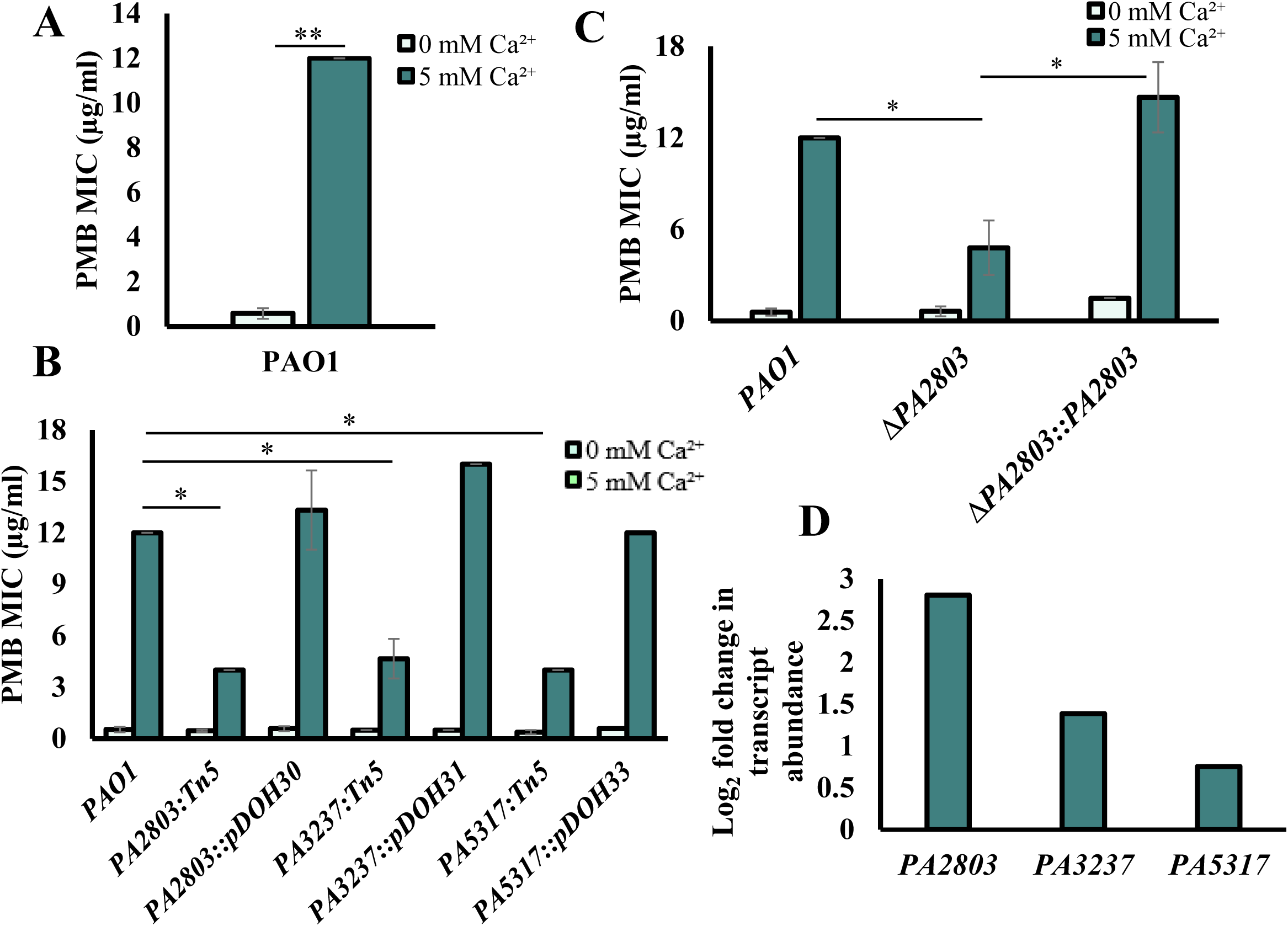
Identifying novel genes involved in Ca^2+^-induced PMB resistance. Light blue; 0 mM Ca^2+^, dark blue; 5 mM Ca^2+^. **(A) PMB resistance in WT PAO1 in response to Ca^2+^.** WT PAO1 was grown without or with 5 mM Ca^2+^, normalized to OD_600_ of 0.1, and plated onto BMM agar plates with the corresponding concentration of Ca^2+^. E-strips with gradient of PMB were placed on the bacterial lawns. MIC was recorded after 24 h incubation at 37⁰C. (**B) E-test for PAO1 and transposon mutants.** Cells were grown, and MIC was determined as in (A). (**C) E-test assay for mutants of PA2803.** WT, Δ*PA2803* and Δ*PA2803*::*PA2803* strains were grown, and MIC was determined as in (A). (**D) RNA-seq analysis**: Log_2_ fold change in transcript abundance of *PA2803*, *PA3237,* and *PA5317* in PAO1 in response to 5 mM Ca^2+^. Student t-test; ** p-value ≤0.01, * p-value ≤0.05.

### Previously identified PMB resistance genes do not contribute to Ca^2+^-dependent PMB resistance in *Pa* PAO1

We first investigated the role of previously identified determinants of *Pa* PMB resistance in Ca²⁺-dependent PMB resistance. The transcriptional profiling of 37 genes reported to contribute to PMB resistance in *Pa* (35–39,92–108) showed that none were induced by Ca²⁺ and some were downregulated in the presence of 5 mM Ca²⁺ (Table 3), including TCSs *phoPQ*, *pmrAB*, *parRS*, *cprRS* and *cbrAB*, and the *arn* operon (109–111). We tested transposon (Tn) mutants with individually disrupted *phoP, pmrB,* and *parR* for Ca²⁺-dependent PMB susceptibility, and found no significant difference compared to the wild-type (WT) strain (Fig. S3A). We also tested a deletion mutant lacking *carR*, encoding the transcriptional regulator of Ca^2+^-responsive TCS CarSR (109) (identical to BqsSR in *Pa* PA14 (112)), and observed no change in PMB MIC in the presence of Ca²⁺. Similarly, deletion of the *mexAB-oprM* operon, encoding the efflux pump implicated in *Pa* polymyxin E resistance (108), did not alter PMB resistance at 5 mM Ca^2+^ (Fig. S3A). These results suggest that Ca²⁺-dependent PMB resistance involves the mechanisms other than the previously characterized determinants.

**Table 3.** Fold change in transcript abundance of known PMB resistance genes in *Pa* PAO1 when grown in the presence of 5 mM Ca^2+^ vs that at no added Ca^2+^. References supporting the role of the selected genes in PMB resistance are provided.

| Mechanism of resistance | Function | PA | Gene name (Ref) | Log2 fold |
| --- | --- | --- | --- | --- |
| **L-Ara4N or PEtn modification of lipid A** | Two-component system (TCS) kinase/response regulator | PA1179 | *phoP* (92,95–97) | -3.8 |
|  |  | PA1180 | *phoQ* | -5.0 |
|  |  | PA4776 | *pmrA*(35,93,98,99) | -3.3 |
|  |  | PA4777 | *pmrB* | -4.1 |
|  |  | PA1799 | *parR*(36) | -0.4 |
|  |  | PA1798 | *parS* | -1.6 |
|  |  | PA3077 | *cprR*(38) | -0.4 |
|  |  | PA3078 | *cprS* | -1.5 |
|  |  | PA4381 | *colR*(100) | 0.4 |
|  |  | PA4380 | *colS* | -1.3 |
|  |  | PA4725 | *cbrA*(39) | -0.3 |
|  |  | PA4726 | *cbrB* | 0.8 |
|  | UDP-glucose dehydrogenase | PA3552 | *arnB* | -6.5 |
|  |  | PA3553 | *arnC* | -6.3 |
|  |  | PA3554 | *arnA* | -5.6 |
|  |  | PA3555 | *arnD*(94) | -6.0 |
|  |  | PA3556 | *arnT* | -5.9 |
|  |  | PA3557 | *arnE* | -6.6 |
|  |  | PA3558 | *arnF* | -8.3 |
| **Repressor of phoPQ expression** | MerR-like | PA4878 | *brlR*(101) | -0.1 |
| **Phosphorylation of lipid A** | Phosphotransfer | PA5009 | *waaP* (*rfaP*) (102) | -0.9 |
| **Surface and membrane remodeling** | Alterations | PA0905 | *rsmA*(103) | -1.8 |
|  | LPS | PA1178 | *oprH*(104) | -6.2 |
| **Efflux and transport** |  | PA2019 | *mexX*(105) | 0.6 |
|  |  | PA2018 | *mexY* | 0.5 |
|  |  | PA0425 | *mexA* | 0.7 |
|  |  | PA0426 | *mexB*(108) | 0.2 |
|  |  | PA0427 | *oprM* | -0.9 |
| **Other polymyxin resistance determinants with either known or unknown functions** |  | PA3533 | *grxD*(106) | -0.1 |
|  |  | PA1588 | *sucC* | -0.1 |
|  |  | PA2023 | *galU* | -0.1 |
|  |  | PA4069 | *PA4069* | 0.1 |
|  |  | PA4109 | *ampR*(107) | 0.2 |
|  |  | PA4459 | *lptC* | -1.1 |
|  |  | PA5000 | *wapR* | -0.5 |
|  |  | PA5001 | *ssg* | -1.2 |
|  |  | PA5199 | *amgS* | -0.3 |

### Three hypothetical proteins of unknown function are involved in Ca^2+^-dependent PMB resistance

To identify genes responsible for Ca^2+^-dependent PMB resistance, we used random chemical mutagenesis followed by PMB susceptibility testing. This led to the isolation of 20 mutant strains with increased sensitivity to PMB in the presence of 5 mM Ca^2+^, compared to WT PAO1. We then complemented four mutant strains with a PAO1 genomic library and isolated colonies that restored the WT level of PMB resistance at 5 mM Ca^2+^ (Fig. S3B). Sequencing the complementing plasmids revealed DNA regions including: *PA2802-PA2803-PA2804*, *PA3237-PA3238*, *PA2590,* and *PA5317*. To characterize the role of the identified genes in Ca^2+^-dependent PMB resistance, the corresponding Tn-mutants were tested for PMB resistance in the presence and absence of Ca^2+^ (Fig. 1B, S3C). The disruption of three genes: *PA2803*, *PA3237* and *PA5317* reduced PMB resistance at elevated Ca^2+^ by more than 50% (Fig. 1B). To verify their role in Ca^2+^-dependent PMB resistance, we complemented the Tn mutants, each with the DNA fragment including the corresponding gene. The complemented strains had restored PMB resistance to the WT level in the presence of Ca^2+^ (Fig. 1B). For further validation, we generated deletion strains, Δ*PA2803, ΔPA3237,* and Δ*PA5317* that also showed a significant reduction in Ca^2+^-dependent PMB resistance (Fig. 1C, S3D). These changes in PMB resistance were not due to mutation-dependent growth defects tested by monitoring growth (Fig. S3E-G). Furthermore, our RNA seq data (PRJNA703087) showed that the transcription of *PA2803*, *PA3237,* and *PA5317* was upregulated at 5 mM vs no added Ca^2+^ by 7, 2.6 and 1.6-fold, respectively (Fig. 1D), further supporting their involvement in Ca^2+^-dependent response.

Sequence analyses predicted that *PA2803*, *PA3237* and *PA5317* encode for a putative phosphonoacetaldehyde hydrolase, a putative DNA binding protein, and a peptide binding component of an ABC transporter, respectively (Fig. S4A and B). The rest of this study is focused on PA2803 and its potential role in Ca^2+^-induced PMB resistance in *Pa*.

### PA2803 encodes for a phosphate and <u>C</u>a^2+^ regulated protein, PcrP

In a genome-wide transcriptional study in *Pa* (24), expression of *PA2803* was shown upregulated by 8.8-fold in response to low P_i_ levels (100 µM vs 1 mM). This regulation is likely mediated by the transcription factor PhoB, known to control P_i_ starvation responses in *Pa* (113) and shown to bind to the promoter region of *PA2803* (Fig. 2A) (114). Supporting these findings, a proteomic analysis of *Pa*’s responses to low P_i_ (50 µM versus 1 mM) also showed the increased abundance of PA2803 by 9.2-fold in response to P_i_ starvation (31), further suggesting that *PA2803* expression is modulated by P_i_ availability.

**Figure 2.**
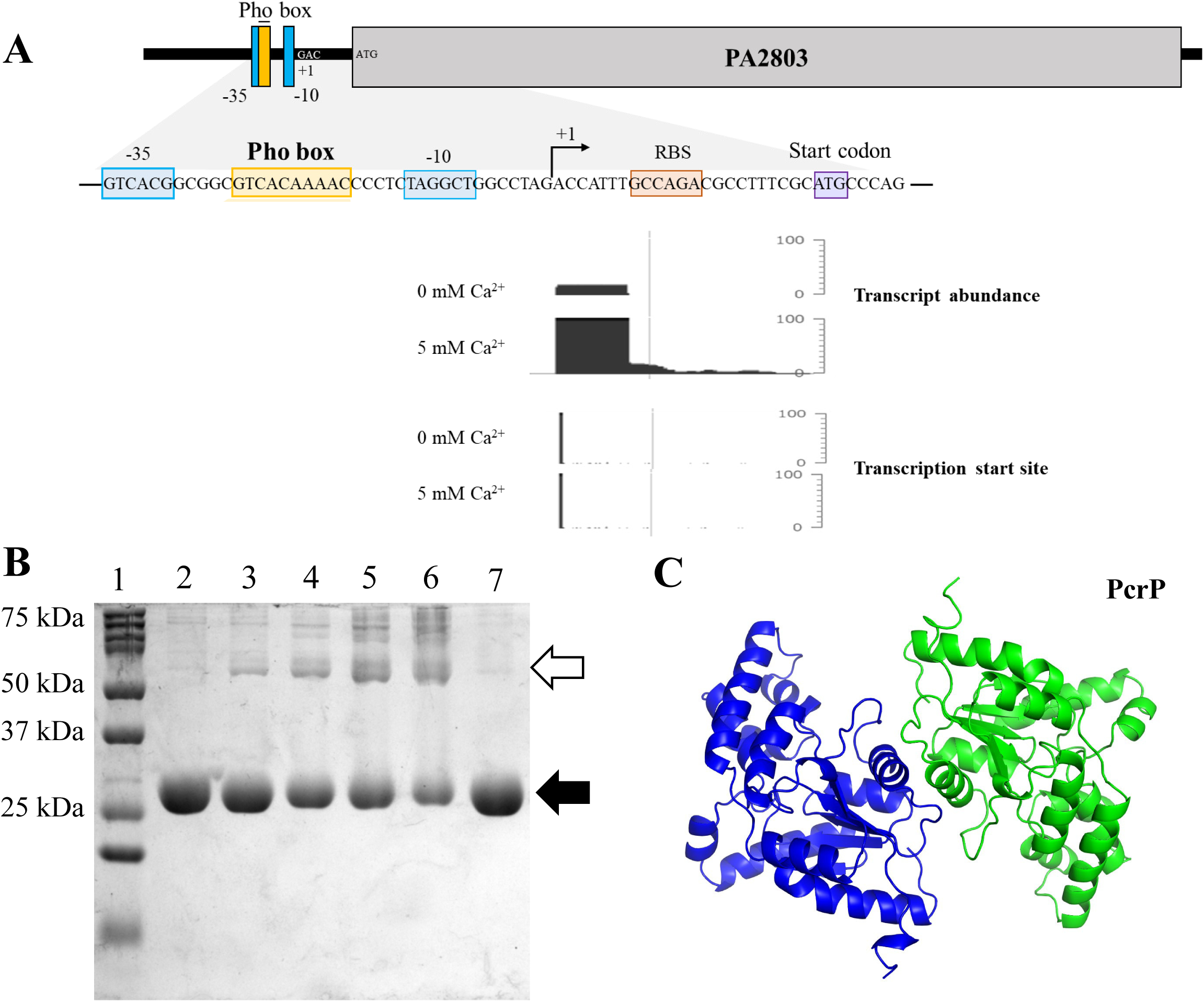
Transcriptional regulation and dimer formation of PA2803. **(A) Promoter region and regulatory elements.** (i) the Pho box region upstream of PA2803 that has been shown to bind PhoB by using EMSA assays (114); (ii) cappable RNA seq transcript abundance and the transcription start site of PA2803 at 0 and 5 mM Ca^2+^. (**B) Oligomerization of PcrP.** Purified PcrP (50 µM) was crosslinked with increasing concentrations of 1% glutaraldehyde and separated on SDS-PAGE gel. Lane 1: protein ladder marker, lane 2: untreated PcrP, lane 3-4, PcrP treated with 0.02%, 0.05%, 0.1% and 0.2% glutaraldehyde. The black and white arrows indicate the bands corresponding to the size of the monomer and dimer of PcrP, respectively. (**C) Molecular dynamics of dimers indicates a stable complex for PcrP**. There two-colored chains indicate two different units in a dimeric complex, at the start of the MD simulations. The grey structures indicate the conformation at the end of 1 μs MD simulation. The 3D structure prediction and *in-silico* dimer simulation of PcrP was performed using Alphafold.

In addition to P_i_ regulation, our RNA seq data (PRJNA703087) revealed that *PA2803* transcription in *Pa* is regulated by Ca²⁺, with a ∼7-fold increase in expression at elevated Ca^2+^ (Fig 1D, 2A). Given this dual regulation, we named the protein encoded by *PA2803,* PcrP, which stands for <u>P</u>_i_ and <u>C</u>a^2+^ <u>r</u>egulated <u>p</u>rotein.

### PcrP provides a Ca^2+^-dependent growth advantage and PMB resistance during P_i_ starvation

Guided by the transcriptomic and proteomic data (24,31), we predicted that PcrP contributes to P_i_ starvation responses in *Pa.* To test its role in *Pa* survival under P_i_ limiting conditions, we monitored growth of PAO1 and Δ*pcrP* in BMMH8 medium supplemented with low (50 µM). The inocula of these strains were pre-grown at no Ca^2+^ and a low P_i_ to prevent accumulation of P_i_ and hindering the impact of P_i_ starvation. The cultures were then inoculated into low or high P_i_ media with or without Ca^2+^. The presence of Ca^2+^ increased the ability of PAO1 to survive P_i_ starvation (Fig. 3A, Fig. S5A). Compared to WT, Δ*pcrP* showed a modest decrease in P_i_-starved growth at 5 mM Ca^2+^ especially during stationary phase that was partially recovered by genetic complementation (Fig. 3A and Fig. S5A). These results, along with our observation of no growth defects in Δ*pcrP* in BMM medium with high (580 µM) P_i_ (Fig. S3E), suggest that PcrP contributes to P_i_ starvation responses in *Pa* in a Ca^2+^-dependent manner. Considering this, we also evaluated whether PcrP plays role in PMB resistance under P_i_ limitation. PMB susceptibility tests on PAO1 and Δ*pcrP* strains were conducted at low (50 µM) P_i_ in the absence or presence of 5 mM Ca^2+^. The results showed that elevated Ca^2+^ enhances PMB resistance in PAO1 at low P_i_ as well, and that about 30% of this resistance is lost in the mutant (Fig. 3B).

**Figure 3.**
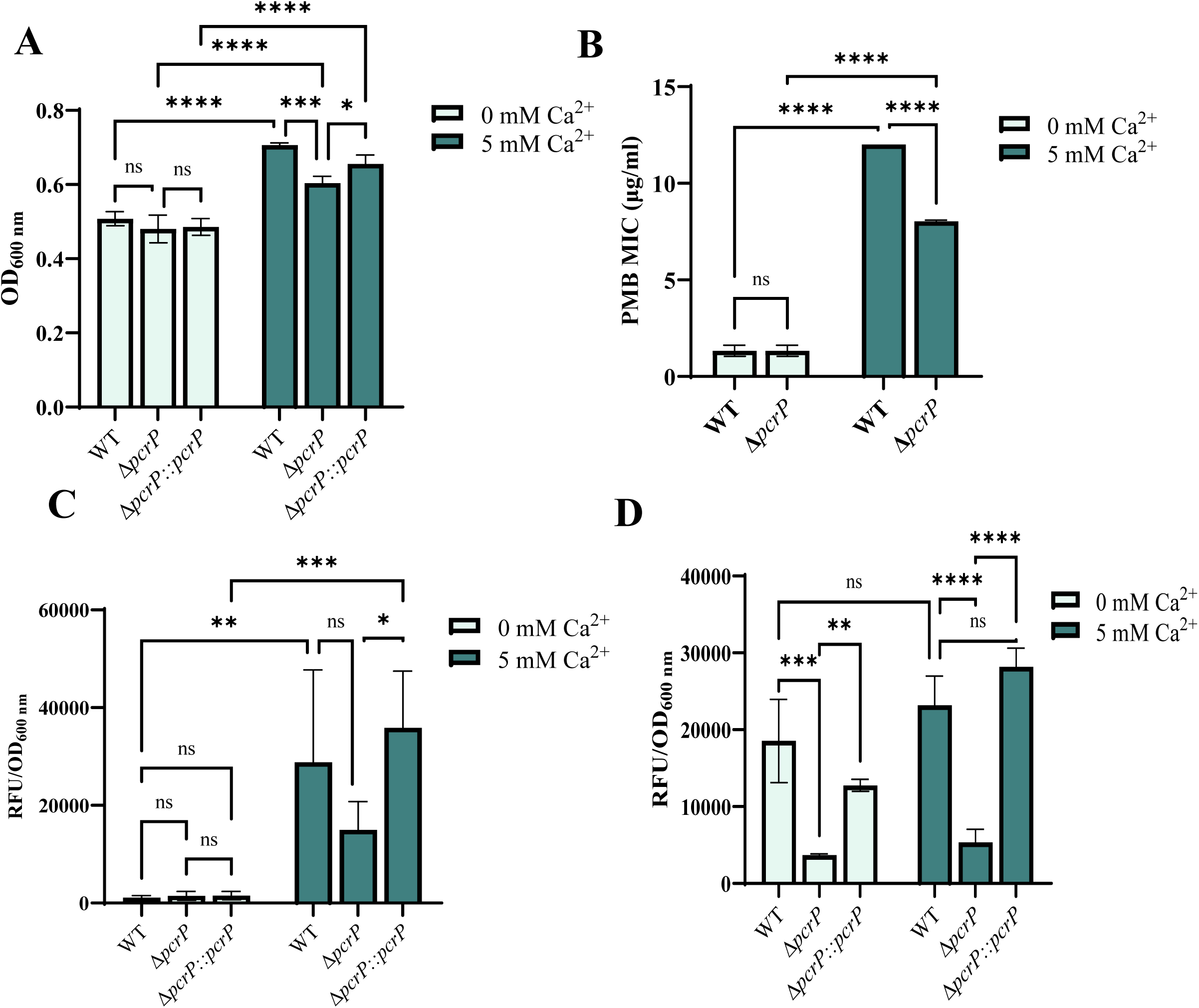
Role of PcrP in P_i_ starvation responses in *P. aeruginosa*. **(A) Growth changes of WT, Δ*pcrP* and Δ*pcrP:*:*pcrP* strains during stationary phase growth at low P_i_.** Cells were grown at limited Pi (50 µM) precultures which were then normalized to OD_600_ of 0.3 and inoculated at 0.1% into low Pi BMMH8 with no or 5 mM Ca^2+^. Growth was monitored for 38 h with OD_600_ reads being taken every 3 h. The growth difference observed 38 h during the stationary phase is shown here. Ca^2+^ enhances the growth of PAO1 and the mutant shows a reduction in growth at 5 mM Ca^2+.^ Light blue; 0 mM Ca^2^, dark blue; 5 mM Ca^2+^. **(B) MIC of PMB at low P_i_ in BMMH8.** Precultures of bacterial strains were grown in BMMH8 supplemented with 50 µM which were then normalized to OD_600_ of 0.1 and spread on low P_i_ BMMH8 agar plates at 0 and −5 mM Ca^2+^. E-test strips carrying a gradient of PMB concentrations were placed on bacterial lawns and the plates were incubated for 24 h at 37⁰C after which the MIC was read. Light blue; 0 mM Ca^2+^, dark blue; 5 mM Ca^2+^. (C) Quantification of poly P levels in WT, Δ*pcrP* and Δ*pcrP::pcrP* strains. Cells grown in high P_i_ (0.58 mM) BMM8 medium at 0-and 5 mM Ca^2+^ for 12 h were collected, washed, normalized to OD_600_ of 0.5 and stained with DAPI for 30 min at room temperature. Fluorescence of the cells were measured at 415/520 nm (ex./em.) and was normalized by OD_600_ to determine the abundance of poly P. Light blue; 0 mM Ca^2+^, dark blue; 5 mM Ca^2+^. Poly P levels in all the strains are elevated in response to Ca^2+^ and the deletion of PcrP shows a reduction comparison to the WT only at 5 mM Ca^2+^. (D) Changes in pyoverdine production in WT, Δ*pcrP* and Δ*pcrP::pcrP* strains during growth at low P_i_. Cells were grown in low Pi (50 uM) media with 0 or 5 mM Ca^2+^ and 200 µl aliquots taken from cultures during growth were used to measure the fluorescence at 400/460 nm (Ex/Em) The fluorescence reads were normalized by OD_600_. Light blue; 0 mM Ca^2+^, dark blue; 5 mM Ca^2+^. Δ*pcrP* shows reduced levels of pyoverdine irrespective of Ca^2+^ concentration. One-way ANOVA; **** p-value <0.0001, *** p-value ≤0.001, ** p-value ≤0.01, * p-value ≤0.05

### PcrP does not impact P_i_ uptake but plays a role in poly P production

To gain further insights, we tested whether the protein is involved in specific P_i_ starvation responses, such as P_i_ uptake, formation of polyphosphates (polyP), and membrane phospholipid homeostasis.

To determine the role of PcrP in P_i_ uptake, PAO1, Δ*pcrP* and Δ*pcrP*::*pcrP* strains were grown at low (50 µM) P_i_ at no added or 5 mM Ca^2+^, and the level of P_i_ in culture supernatants was monitored using malachite green assay. Although a slight decrease in P_i_ uptake was observed at 5 mM Ca^2+^, this response was not due to the deletion of *pcrP* (Fig. S5B), indicating that PcrP does not play a role in P_i_ uptake of *Pa* under the conditions tested.

Next, we evaluated the role of PcrP in the formation of poly P complexes in *Pa*. Poly P complexes are made of linear polymeric chains of P_i_ residues ranging from 3 to more than 1000, linked by high-energy phosphoanhydride bonds (115,116). We hypothesized that the involvement of PcrP in poly P formation may explain the observed growth defect in the mutant. To test this, PAO1 and Δ*pcrP* strains were grown in BMM8 medium with high P_i_ (580 µM, to support accumulation of Poly P) at no added or 5 mM Ca^2+^ and followed by Hoechst staining. We observed a 25-fold Ca^2+^-dependent increase in the poly P levels in PAO1 with a 48 % reduction in Δ*pcrP* strain at 5 mM (Fig. 3C). The mutant poly P level was restored to the WT levels by gene complementation (Fig. 3C). This suggests that elevated Ca^2+^ enhances the accumulation of poly P in *Pa* and that PcrP plays a role in this process.

Since membrane phospholipid remodeling occurs during *Pa* adaptation to P_i_ starvation (14), we evaluated possible involvement of PcrP in this process. We were particularly interested in the production of membrane lipids with non-phosphate-containing head groups, such as ornithine lipids (OL), which is a well-established mechanism of adapting to limiting P_i_ (14). To assay the membrane OL content, WT and Δ*pcrP* strains were grown at low (50 µM) P_i_ conditions and their membrane lipids extracts were separated by TLC followed by ninhydrin-based detection for amino-lipids. We detected lipid spots indicative of the presence of amino lipids, such as OL, under P_i_ limiting conditions (Fig. S5C), consistent with previous reports (14). An increase in the intensity of these lipid spots was apparent at 5 mM Ca^2+^ in WT, while their abundance appeared greater in the Δ*pcrP* strain, indicating Ca²⁺-dependent changes in the abundance of these lipids in *Pa* and PcrP role in their biosynthesis.

### PcrP plays a role in the production of virulence factors, pyocyanin and pyoverdine under P_i_ starvation

P_i_ limitation is the condition that bacteria frequently face during infections (26), and many virulence factors in *Pa* are regulated by P_i_ availability, including toxin pyocyanin and siderophore pyoverdine (26,29). Based on PcrP role in *Pa* responses to P_i_ limitation, we hypothesized that the protein may also have an impact on P_i_ regulation of *Pa* virulence. Although only a minor decrease of 16-18% in the pyocyanin level was observed in the mutant at low P_i_ (Fig. S5D), 77-80% decrease in pyoverdine level was detected at both Ca^2+^ levels (Fig. 3D). These findings indicate that PcrP contributes to shaping the *Pa*’s virulence factor production during P_i_ starvation.

### PcrP is the founding member of the “PA2803 subfamily” of Haloacid Dehalogenase Superfamily HADSF

Sequence analyses and protein structure modeling showed that PcrP carries a conserved α/β-domain classified as a hydrolase fold that is shared among a broad superfamily of Haloacid dehalogenase superfamily (HADSF) proteins (Interproscan ID: IPR023214). This superfamily includes approximately one million members (Interpro database) that vary widely in their taxonomical distribution, domain architecture, and biological functions (46). Members are present in more than 40,000 species, spanning all three domains of life. The genus of *Pseudomonas* encodes about 485 HADSF proteins that are spread among seven species (Fig. S1). In addition to PcrP, the PAO1 genome encodes another member of the HADSF, PhnX, that shares 27.4% amino acid identity with PcrP and has an established function of 2-phosphonoacetaldehyde hydrolase (117–120). According to the earlier reports (24) and our RNA-seq data (log_2_ fold-change of 0.42 for *phnX* at 5 mM vs 0 mM Ca^2+^), unlike PcrP, PhnX is not regulated by P_i_ and Ca^2+^.

Phylogenetic analysis of PcrP and its closest 12 homologs, including PhnX from PAO1 and PSPTO_2114 from *P. syringae* (49), showed that PcrP and PSPTO_2114 form a distinct clade within C1-cap containing phosphonatases (Fig. 4A), separating them from functionally characterized phosphonatases (Fig. 4B). This is consistent with the report in (49), designating the subfamily of *Pseudomonas* proteins that cluster with PA2803 as the “PA2803 subfamily”. PA2803 lacks the conservation of catalytically active residues in functional phosphonatases (nucleophilic Asp and Thr in motif 1, Lys in cap domain (absent), Ser/Thr in motif 2, Lys/Arg in motif 3, and two Asp/Glu residues in motif 4) and contains an abbreviated cap domain (Fig. 4C). It is noteworthy that among the identified members of the PA2803 subfamily, PcrP retains a larger portion of the cap domain in comparison to its homolog PSPTO_2114 that was found catalytically inactive (49).

**Figure 4.**
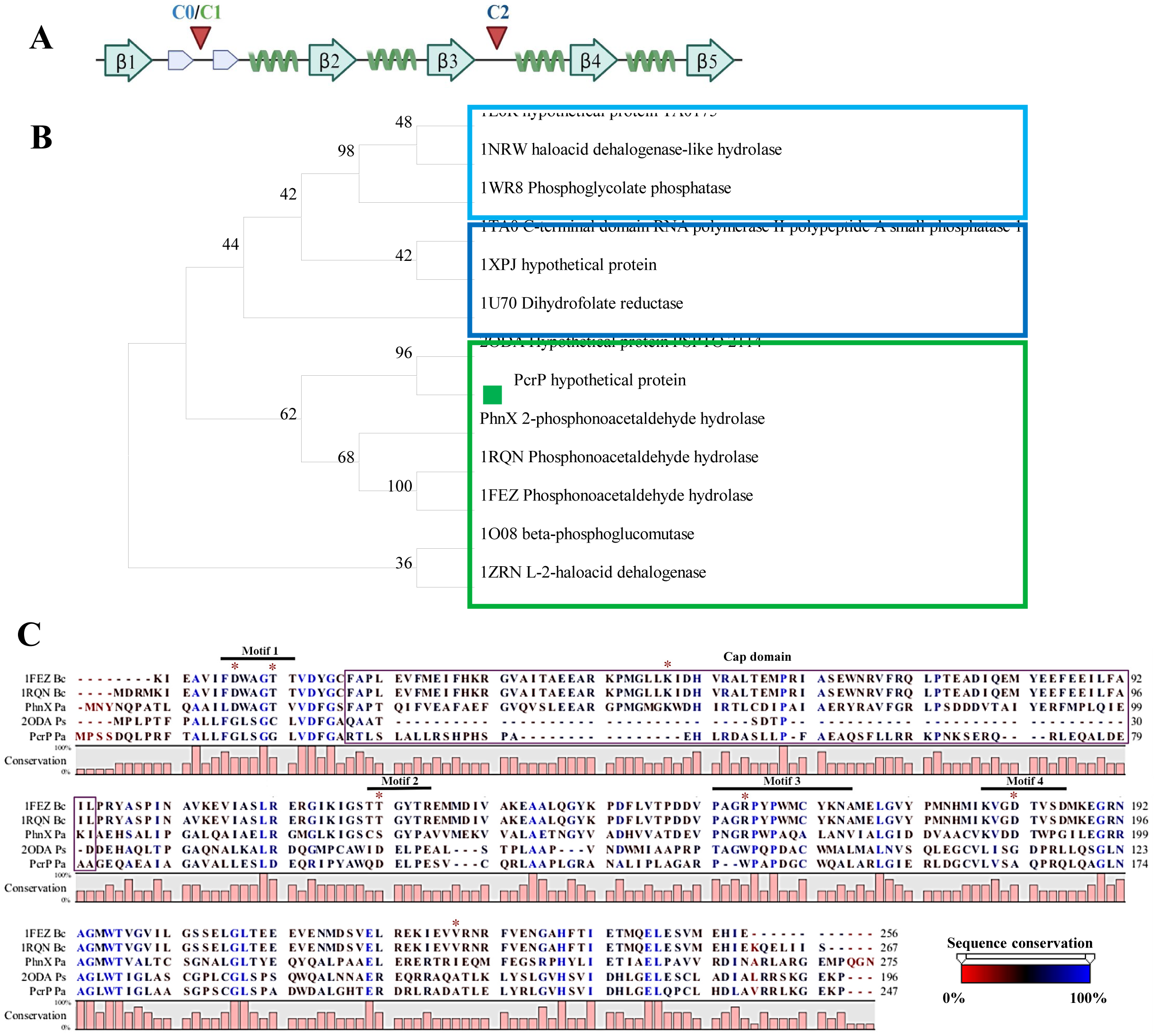
Position of the CAP domain within HADSF proteins and sequence conservation and evolutionary relationships of PcrP. **(A)** Typical haloacid dehalogenase (HAD) superfamily’s core domain is an α/β sandwich with a characteristic Rossmannoid α/β fold structure. In this linear topology, β-strands and alpha-helices are represented with blue arrows and green helices, respectively. β-Strands are numbered 1-5. Small blue arrows located between β1 and β2 represent the β-hairpin motif referred to as flap, a key structural feature of HAD domain crucial for catalytic function. Red arrows indicate the two different positions where the cap domains are inserted within different HADSF proteins, with C0 and C1 caps inserted between the two β-strands of the flap and C2 caps inserted immediately after the β3 strand of the core domain**. (B) Phylogenetic relationships of PcrP inferred by maximum likelihood tree.** PcrP amino acid sequence was aligned to its paralog PhnX and 12 other proteins that are representative of different C0, C1 and C2 cap domains described in Burroughs et al., 2006 (42). Sequence identifiers indicate their PDB code/Protein name followed by predicted function. The alignment was done using MUSCLE and the evolutionary history was inferred by using the Maximum Likelihood method and JTT matrix-based model (163) in MEGA11 (164). The bootstrap values inferred from a bootstrap consensus tree of 50 replicates are shown next to the branches. Clades that separate C0, C1 and C2 are represented within squares that correspond to their cap domain architectures as depicted in (**A**). PcrP, denoted by a green square and its closest homolog PSPTO_2114 (PDB ID:2ODA) forms a separate clade within the C1 group. (**C) Multiple sequence alignment displaying the catalytic residues within PcrP in comparison to phosphonatases.** Proteins containing a C1-type cap domain highlighted within the bracket (PhnX from *B. cereus:* 1FEZ and 1RQN, PhnX from *Pa* PAO1, PSPTO_2114 from *P. syringae pv. tomato* and PcrP) were aligned using MUSCLE. The bar graph indicates the % conservation and amino acid positions placed above the alignment in relation to the sequence of PcrP. Residues are colored based on the sequence conservation with red to blue representing the range 0-100%. C1 cap domain found in phosphonatases is depicted in purple and is truncated in PcrP as indicated. Amino acid residues governing catalytic activity and cofactor binding identified in the functionally characterized phosphonatase 1FEZ from *Bacillus cereus* are highlighted with red asterisks. PcrP and its homolog PSPTO_2114 lacks conservation of these residues implying the loss of phosphonatase activity.

### PcrP shows no catalytic activity

To assess whether PcrP has any catalytic activity, we evaluated its phosphonatase and phosphatase activities using phosphonoacetaldehyde and para-nitro phenol phosphate (pNPP) as substrates, respectively. For this, PcrP was expressed and purified from *E. coli* BL21. As predicted by sequence analysis, we did not detect any phosphonatase nor phosphatase activity (Fig. S6A and B), suggesting that in spite of the larger portion of the cap domain, similarly to its homolog PSPTO_2114 (49), PcrP lacks catalytic activity.

Since functionally active phosphonatases are generally known to form dimers (49,121), as exemplified by the phosphonatase from *Bacillus cereus* (PDB ID:1FEZ) (47), this ability may play an important role in their enzymatic function. Therefore, we aimed to determine whether the lack of catalytic activity in PcrP is related to its inability to dimerize. However, upon crosslinking the recombinant PcrP with increasing concentrations of glutaraldehyde, we observed a protein band with the molecular weight corresponding to PcrP dimer (Fig. 2B). This verifies that the oligomeric state of PcrP is not the cause of the catalytic activity loss.

### PcrP binds protein partners, including PA3518 and Acp3

It was previously suggested that the PA2803 subfamily proteins may be able to interact with other proteins (49). Therefore, we investigated possible protein-protein interactions of PcrP. For this, Ni affinity-based pull-down and Flag-based co-immunoprecipitation assays were performed by using 6X-His-tagged PcrP and 3X-Flag-tagged PcrP expressed in *Pa* Δ*pcrP* grown at 0 or 5 mM Ca^2+^. As a negative control, *Pa* Δ*pcrP* lacking the native PcrP was used. For protein identification, we considered both LC-MS/MS confidence score (q-value <0.01), consistency of the protein detection by both 6X-His and 3X-Flag-based approaches, and cellular localization. Our analysis identified 202 putative binding partners of PcrP. This list was curated to include the proteins whose abundance increased by a log_2_ fold-change of 5 compared to that in the Δ*pcrP* mutant and reached at least 25% of the bait (PcrP) abundance under identical conditions, thereby minimizing the potential for false-positive hits. This resulted in 22 potential interacting partners of PcrP (Table 4). The top three hits, PqsB, PA3518, and Acp3, were chosen for validation using BTH assays. In these assays, PcrP served as the bait, while each of the selected putative binding partners acted as the prey. We evaluated their interactions both qualitatively by blue/white colony screening and quantitatively by using B-gal assay. Since we PcrP formed dimers during crosslinking (Fig. 2B), we used BTH to confirm PcrP dimerization in cells (Fig. 5A) using T18-PcrP and T25-PcrP fusions. As a positive control, we used *E. coli* cells carrying T25 and T18 fragments fused to the established leucine-zipper dimerization motif (68). As negative controls, we used colonies carrying a pair of empty vectors pUT18C and pKT25 and the pairs of the constructs carrying T18-PcrP (or other tested proteins) with its empty vector counterpart (e.g., pUT18C:*pcrP* and pKT25). Out of the tested putative partners, PA3518 and Acp3 were verified to interact with PcrP (Fig. 5A).

**Figure 5.**
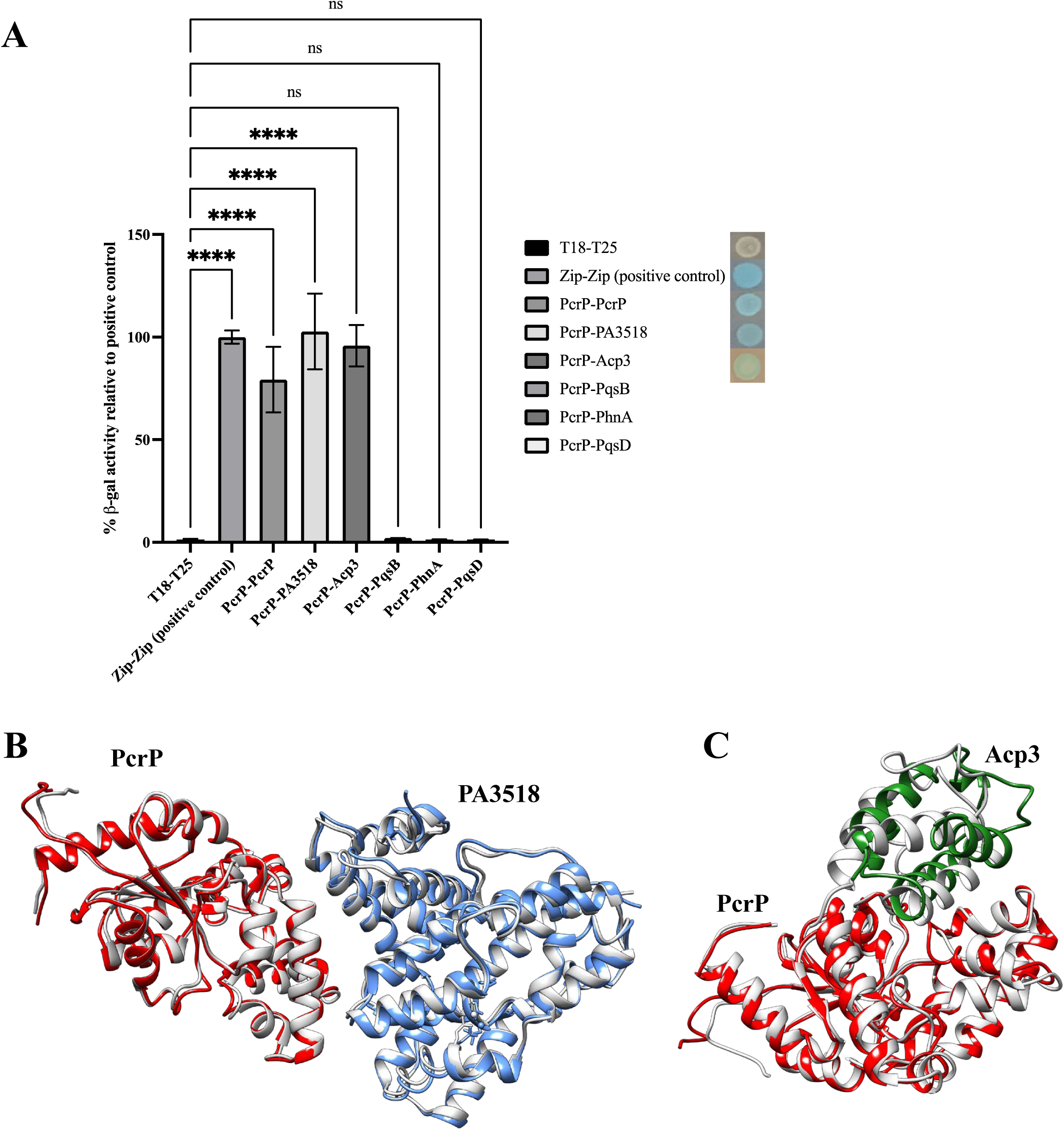
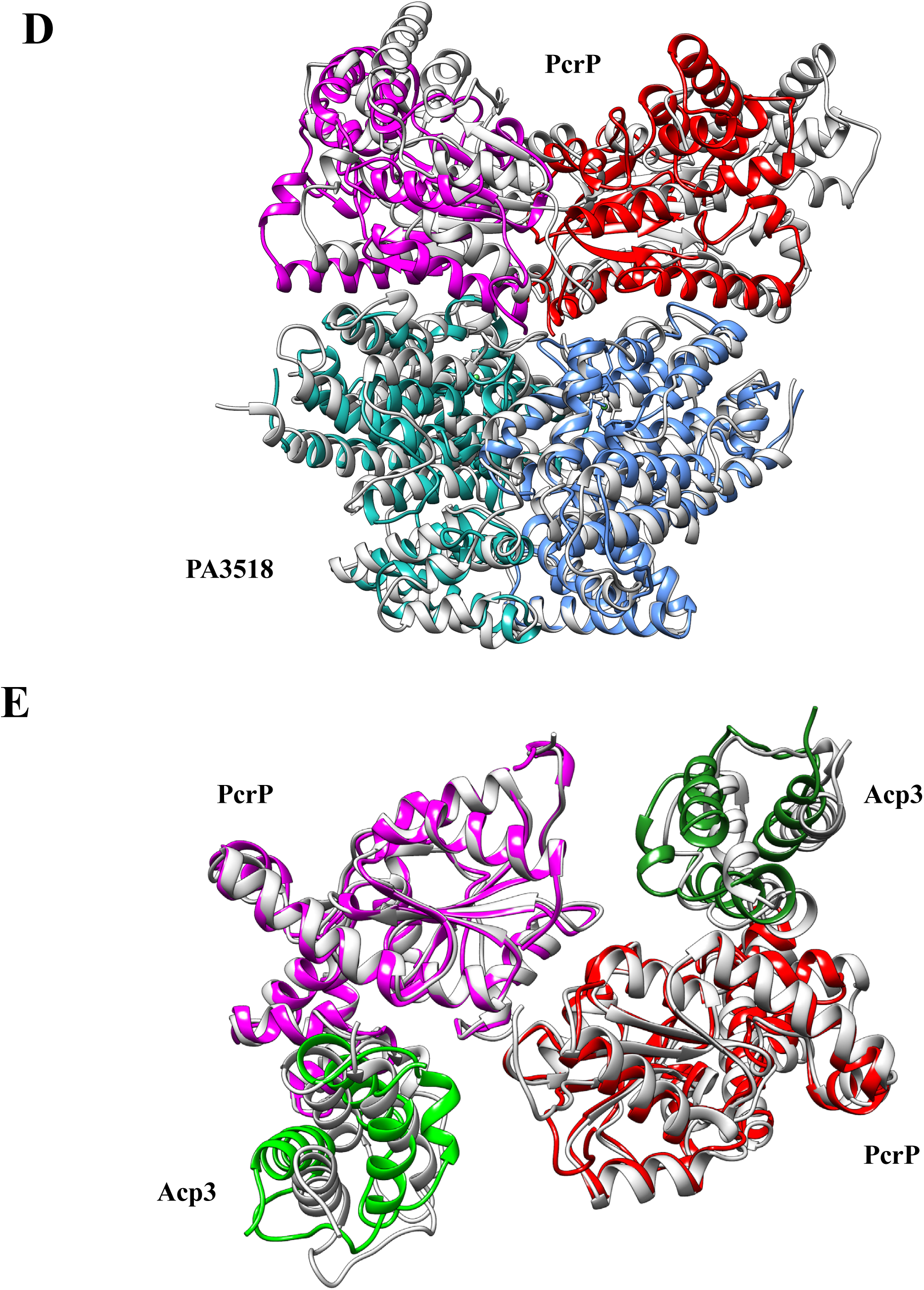
Protein binding function of PcrP. **(A) Bacterial two hybrid assays validating protein partners of PcrP.** Putative partners of PcrP identified among a curated list of proteins were cloned into pUT18C vector as bait, and *pcrP* into pKT25 as prey. The plasmid pairs were co-transformed into subjected to *E. coli* BTH101 and subjected to bacterial two hybrid assays. The graph indicates the relative β-gal activity during protein interaction with their respective blue-white colony formation represented next to the legend. One-way ANOVA: **** p-value <0.0001 **Molecular dynamics of hetero-dimers indicate stable complexes of PcrP-PA3518 (B) and Pcrp-Acp3** (**C**). The two-colored chains indicate two different units in a hetero-dimeric complex, at the start of the MD simulations. The grey structures indicate the conformation at the end of 1 μs MD simulation. **Molecular dynamics of hetero-tetramer indicate stable complexes of PcrP-PA3518 complex (D) and Pcrp-Acp3 complex** (**E**). The colored chains indicate conformation at the start of the MD simulations. The grey structures indicate the conformation at the end of 1 μs MD simulation.

**Table 4.**
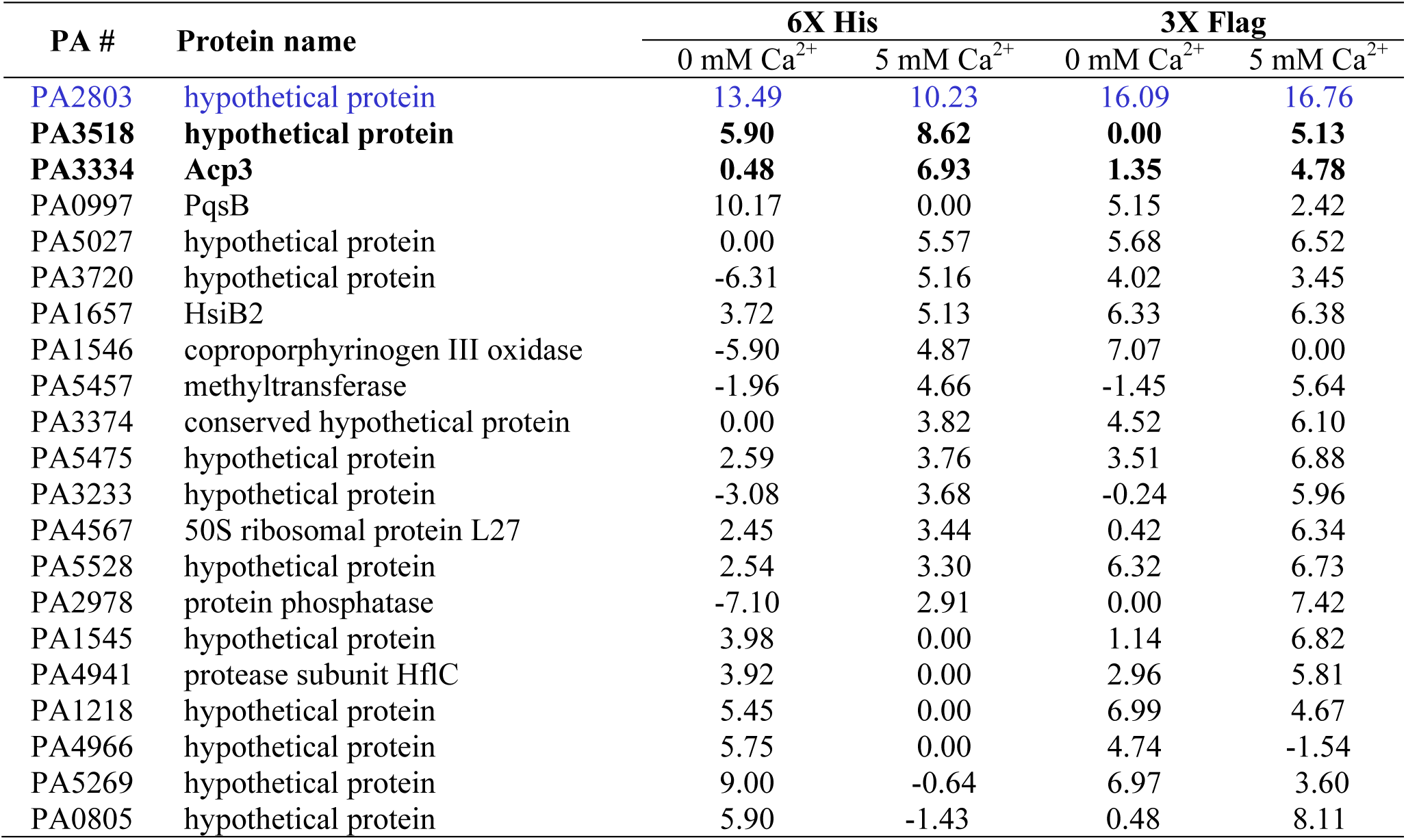
Putative protein binding partners of PcrP identified by pull-down and immunoprecipitation assays using PAO1 strains overexpressing 6XHis and 3XFlag tagged PcrP, respectively. Cells were grown at 0 mM Ca^2+^ or 5 mM Ca^2+^, and Δ*pcrP* was used as a negative control. The values are presented as log_2_ values of the ratio of intensity (tagged-PcrP/Δ*pcrP*). PcrP is highlighted in blue and its protein partners with validated interactions are highlighted in bold.

| PA # | Protein name | 6X His |  | 3X Flag |  |
| --- | --- | --- | --- | --- | --- |
|  |  | 0 mM Ca<sup>2+</sup> | 5 mM Ca<sup>2+</sup> | 0 mM Ca<sup>2+</sup> | 5 mM Ca<sup>2+</sup> |
| PA2803 | hypothetical protein | 13.49 | 10.23 | 16.09 | 16.76 |
| PA3518 | hypothetical protein | 5.90 | 8.62 | 0.00 | 5.13 |
| PA3334 | Acp3 | 0.48 | 6.93 | 1.35 | 4.78 |
| PA0997 | PqsB | 10.17 | 0.00 | 5.15 | 2.42 |
| PA5027 | hypothetical protein | 0.00 | 5.57 | 5.68 | 6.52 |
| PA3720 | hypothetical protein | -6.31 | 5.16 | 4.02 | 3.45 |
| PA1657 | HsiB2 | 3.72 | 5.13 | 6.33 | 6.38 |
| PA1546 | coproporphyrinogen III oxidase | -5.90 | 4.87 | 7.07 | 0.00 |
| PA5457 | methyltransferase | -1.96 | 4.66 | -1.45 | 5.64 |
| PA3374 | conserved hypothetical protein | 0.00 | 3.82 | 4.52 | 6.10 |
| PA5475 | hypothetical protein | 2.59 | 3.76 | 3.51 | 6.88 |
| PA3233 | hypothetical protein | -3.08 | 3.68 | -0.24 | 5.96 |
| PA4567 | 50S ribosomal protein L27 | 2.45 | 3.44 | 0.42 | 6.34 |
| PA5528 | hypothetical protein | 2.54 | 3.30 | 6.32 | 6.73 |
| PA2978 | protein phosphatase | -7.10 | 2.91 | 0.00 | 7.42 |
| PA1545 | hypothetical protein | 3.98 | 0.00 | 1.14 | 6.82 |
| PA4941 | protease subunit HflC | 3.92 | 0.00 | 2.96 | 5.81 |
| PA1218 | hypothetical protein | 5.45 | 0.00 | 6.99 | 4.67 |
| PA4966 | hypothetical protein | 5.75 | 0.00 | 4.74 | -1.54 |
| PA5269 | hypothetical protein | 9.00 | -0.64 | 6.97 | 3.60 |
| PA0805 | hypothetical protein | 5.90 | -1.43 | 0.48 | 8.11 |

Given that in addition to PqsB, several other proteins involved in PQS biosynthesis were detected (Table 4), we also evaluated the interaction between PcrP and PqsD and PhnA. However. no β-gal activity was detected (Fig. 5A), which either rules out these interactions or suggests their transient nature or the involvement of other *Pa* proteins.

### PcrP binds to distinct protein partners under P_i_ starvation

Based on our discovery that PcrP binds protein partners, we hypothesized that PcrP may engage in different protein-protein interactions under low P_i_ conditions, which would enable its role in P_i_ starvation responses. To explore this possibility, we performed co-immunoprecipitation assays with *Pa* cell lysates overexpressing 3xFlag-PcrP grown under low P_i_ at no or 5 mM Ca^2+^. To avoid the bias introduced by the difference in the abundance of PcrP at 5 mM Ca^2+^, we focused on the proteins that were enriched by more than 25% of the abundance of the bait at elevated Ca^2+^ concentration. Our analysis revealed six proteins that were detected as putative binding partners at both high and low P_i_ levels, and 17 that were only detected at low P_i_, including HisI (PA5066), PurF (PA3108) and RpiA (PA0330) (Table 5). Although pending validation, these low P_i_-specific protein partners suggest a distinct role of PcrP during phosphate starvation conditions, commonly present in the host environment (26,31).

**Table 5.** Putative protein binding partners of PcrP identified by immunoprecipitation using strains overexpressing 3XFlag tagged PcrP at low P_i_ (50 µM) and high P_i_ (580 µM). Δ*pcrP* strain was used as a negative control. The values are represented as log_2_ values of the ratio of intensity (tagged-PcrP/Δ*pcrP*). Proteins that were enriched by more than 25% of the bait (PcrP) at their respective condition are listed. The top panel represents proteins that were identified at both P_i_ levels, whereas the bottom panel indicates proteins only identified at low P_i_.

| PA # | Protein name | Low $P_i$ | | High $P_i$ | |
| --- | --- | --- | --- | --- | --- |
| | | 0 mM $Ca^{2+}$ | 5 mM $Ca^{2+}$ | 0 mM $Ca^{2+}$ | 5 mM $Ca^{2+}$ |
| PA2803 | PcrP | 6.92 | 10.59 | 16.09 | 16.76 |
| PA1681 | chorismate synthase | ND | 5.43 | 6.50 | 6.53 |
| PA3729 | conserved hypothetical protein | ND | 4.97 | 1.79 | 6.99 |
| PA5015 | pyruvate dehydrogenase | ND | 4.73 | 1.65 | 4.74 |
| PA3831 | leucine aminopeptidase | ND | 4.68 | ND | 5.12 |
| PA4241 | 30S ribosomal protein S13 | ND | 4.12 | ND | 7.36 |
| PA5028 | conserved hypothetical protein | ND | 4.04 | 4.80 | 1.58 |
| PA3654 | uridylate kinase | ND | 3.03 |  | 8.05 |
| PA4352 | conserved hypothetical protein | ND | 3.00 | 1.25 | 7.27 |
| PA4992 | hypothetical protein | ND | 2.70 | 1.41 | 4.61 |
| PA3168 | DNA gyrase subunit A | 2.33 | ND | ND | 6.08 |
| PA5066 | phosphoribosyl-AMP cyclohydrolase | ND | 8.21 | ND | ND |
| PA3108 | amidophosphoribosyltransferase | ND | 7.69 | ND | ND |
| PA0330 | ribose 5-phosphate isomerase | ND | 6.96 | ND | ND |
| PA1585 | 2-oxoglutarate dehydrogenase (E1 subunit) | ND | 4.85 | ND | ND |
| PA4026 | probable acetyltransferase | ND | 4.37 | ND | ND |
| PA3645 | (3R)-hydroxymyristoyl-[acyl carrier protein] dehydratase | ND | 4.28 | ND | ND |
| PA3155 | UDP-2-acetamido-2-dideoxy-d-ribo-hex-3-uluronic acid transaminase, wbpE | ND | 4.20 | ND | ND |
| PA1833 | probable oxidoreductase | ND | 3.90 | ND | ND |
| PA2189 | hypothetical protein | ND | 3.80 | ND | ND |
| PA2173 | hypothetical protein | 4.22 | 3.70 | ND | ND |
| PA4518 | hypothetical protein | ND | 3.64 | ND | ND |
| PA3712 | hypothetical protein | ND | 3.63 | ND | ND |
| PA0542 | conserved hypothetical protein | ND | 3.48 | ND | ND |
| PA0945 | phosphoribosylaminoimidazole synthetase | ND | 2.98 | ND | ND |
| PA3479 | rhamnosyltransferase chain A | 2.26 | 0.00 | ND | ND |
| PA5049 | 50S ribosomal protein L31 | 3.72 | 0.00 | ND | ND |
| PA4671 | probable ribosomal protein L25 | 2.33 | 0.00 | ND | ND |

### Computer modeling supports interactions between PcrP and its partners: PA3518 and Acp3

To generate further insights into the interaction between PA3518 and Acp3 with PcrP, we performed computer simulations using molecular dynamics (MD) based on the Alphafold predicted structure of PcrP (Fig. S4B) and the available X-crystal structures of PA3518 (PDB ID: 3BJD) and solution NMR structure of Acp3 (PDB ID: 2LTE). For a thorough analysis, several different sets of complexes were modeled that consisted of (i) homodimers for PcrP, PA3518 and Acp3; (ii) hetero-dimers PcrP-PA3518 and PcrP-Acp3; (iii) hetero-tetramer complexes of PcrP with PA3518 and separately with Acp3 (both simulations modeled as dimer of dimers); and (iv) hetero-hexamer complex of PcrP with PA3518 and Acp3. Supporting our cross-linking, pull-down, and BTH data, the results indicate that PcrP forms homodimers (Fig. 2C) that are stable during 1 μs MD simulations. Stability in MD simulations is defined as no significant changes from starting structure and the complex retaining majority of the initial inter-chain/inter-protein interactions. Similarly, PA3518 (Fig. S7A) and Acp3 (Fig. S7B) also showed the formation of stable homodimers during 1 μs MD simulations. Further supporting the pull-down and BTH data, MD showed the formation of stable hetero-dimeric complexes between PcrP and PA3518 (Fig. 5B) as well as PcrP and Acp3 (Fig. 5C). We also modeled a possible formation of dimer of dimers, and both the PcrP-PA3518 (Fig. 5D) and PcrP-Acp3 (Fig. 5E) complexes were stable during the MD simulations, supporting the interactions. Finally, we modeled the possibility of the simultaneous interaction of all three proteins using either Alphafold-predicted or manually simulated complex of PcrP-PA3518-Acp3. Interestingly, the Alphafold predicted complex structure showed Acp3 only interacting with PA3518, but not PcrP (Fig. S7C). The alternative approach based on the manually curated two hetero-tetramer complexes of PcrP-PA3518 and PcrP-Acp3 resulted in stable interaction between Acp3 and PcrP at the original binding site of the latter (Fig. S7D) during the entire 1 μs MD simulation. Overall, the computer modeling supported that both PA3518 and Acp3 interact with PcrP individually and indicated that the proteins may interact simultaneously.

In addition, we performed an interaction energy analysis that revealed several potentially insightful observations. For this, the conformations collected during the MD simulations were used to calculate interaction energy between the individual protein chains of the complexes as a sum of electrostatic and van der Waals energy. The results, depicted in Figure S8, support that PcrP forms stable interactions in a homodimer and with Acp3 in a heterodimer (Fig. S8A, where the large negative values indicate strong interaction). PcrP-PcrP interaction in its homodimer form appears to be stronger (average interaction value about −1.8 kcal/mol/residue) compared to the same (interchain) interactions in the hetero-tetramer (PcrP-PA3518, about −1.0 kcal/mol/residue) or the hexa-tetrameric PcrP-PA3518-Acp3 complexes (about −0.5 kcal/mol/residue). Comparatively, PcrP interaction with PA3518 appears to be weaker in the hetero-tetramer (about −1.5 kcal/mol/residue) and hexameric (about −0.9 kcal/mol/residue) complexes (Fig. S8B). The hetero-dimeric complex (PcrP-PA3518) shows increasing interactions over the course of the MD simulation (black line), but the energy level is highly variable, possibly indicating a lack of a clear binding site. As indicated by the largest negative values, the interaction of PcrP with Acp3 (Fig. S8C) appeared very stable in the dimeric, tetrameric, and hexameric complexes. Overall, the energy analyses suggested that PcrP forms stable interaction with Acp3 (indicated by greater and more stable negative values, Fig. S8C), while its interactions with PA3518 may have a more transient nature (smaller negative values with some variations over time, Fig. S8B). An interesting observation is that the PcrP-Acp3 shows more stable interaction in the hexameric (PcrP-PA3518-Acp3) complex compared to tetrameric complex (PcrP-Acp3) or even its dimeric complex (PcrP-Acp3), supporting that the three proteins may form a complex simultaneously.

### PcrP interaction with Acp3 may play a role in *Pa* oxidative stress

Acp3, encoded by PA3334, is one of the three acyl carrier proteins (ACPs) produced by *Pa* (AcpP, Acp1, and Acp3). This protein has been linked to oxidative stress responses in *Pa,* as its deletion resulted in increased resistance to hydrogen peroxide (122). Recent studies showed that Acp3 binds to and represses the major catalase KatA, leading to increased accumulation of reactive oxygen species (ROS) in *Pa* (122) (Fig. 6A). Since PMB antimicrobial effect in *Pa* is mediated by ROS accumulation (123), we hypothesized that PcrP may protect *Pa* from PMB exerted oxidative stress by interacting with Acp3 and alleviating KatA repression. Accordingly, we predicted that upon PcrP-Acp3 binding at high Ca^2+^, ROS accumulation is decreased (Fig. 6A). To test this, we measured the endogenous ROS levels in PAO1 and Δ*pcrP* strains grown with or without Ca^2+^ using the ROS-sensitive dye H_2_DFFA. Consistent with our hypothesis, we observed a 77% decrease in the endogenous ROS levels in the WT PAO1 strain in response to Ca^2+^ (Fig. 6B). This Ca^2+^-dependent release of oxidative stress may provide a protection against PMB and explain the observed Ca^2+^-enhanced PMB resistance. The Δ*pcrP* strain accumulated nearly 2-fold more ROS than WT at 5 mM Ca^2+^ (Fig. 6B) with no difference at no Ca^2+^, suggesting a modest contribution of PcrP to the protective effect of Ca^2+^ against endogenous oxidative stress and, therefore, PMB.

**Figure 6.**
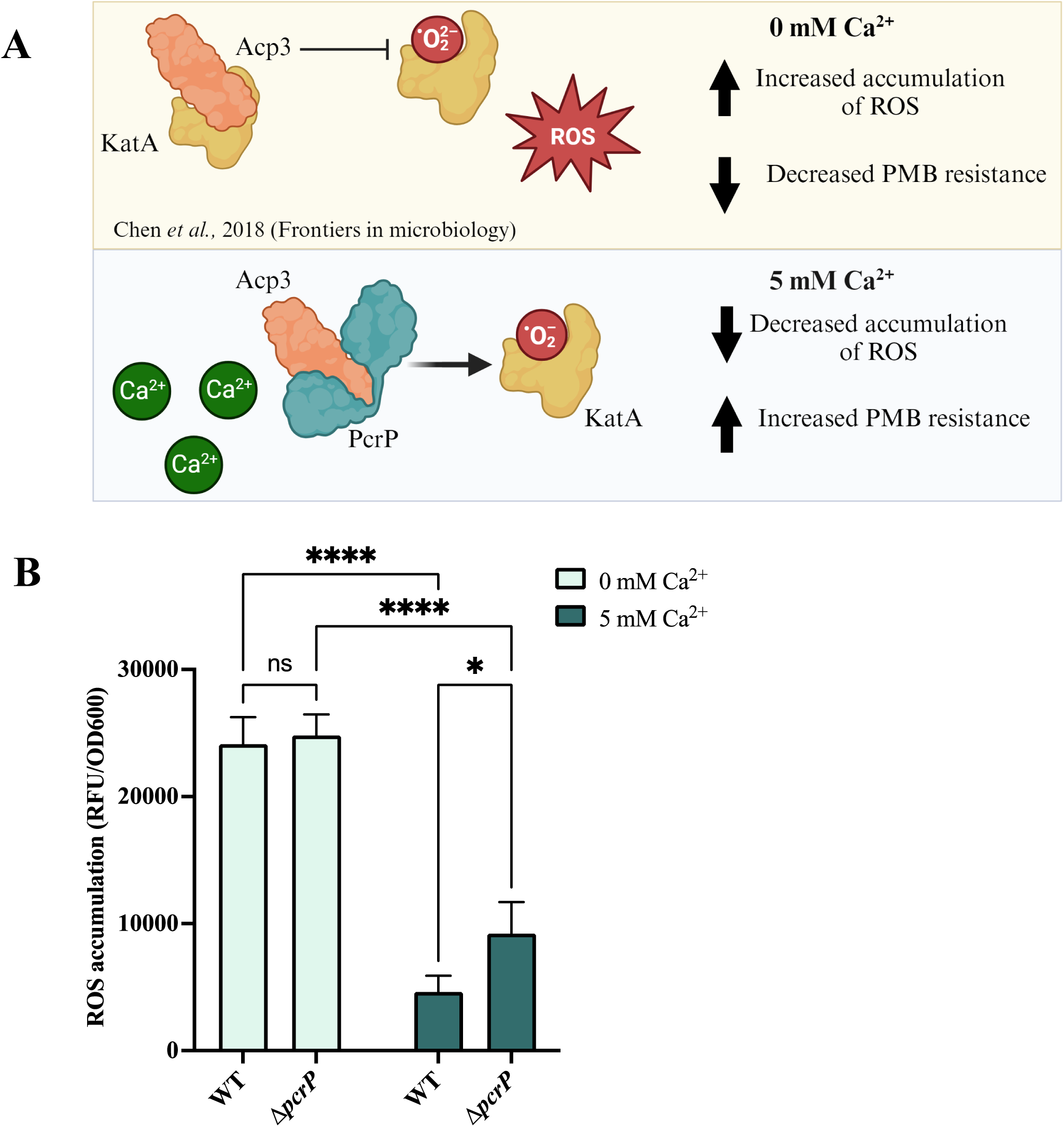
Model illustrating the interaction of Acp3 with PcrP and its impact on oxidative stress in *Pa*. **(A) Current model of PcrP-Acp3 interaction.** Acp3 (acyl carrier protein 3, encoded by PA3334), binds to and inhibits the catalase KatA, leading to increased ROS accumulation (122). Under high Ca^2+^ conditions, PcrP interacts with Acp3, potentially relieving KatA inhibition, thereby reducing oxidative stress. This model predicts that in WT *Pa* with 5 mM Ca^2+^, there will be reduced ROS levels, enhancing PMB resistance. Conversely, a *Pa* strain lacking PcrP at 5 mM Ca^2+^ is expected to exhibit elevated ROS accumulation. (**B) Elevated Ca^2+^ reduces ROS accumulation in WT and Δ*pcrP Pa.*** Fluorescence of the ROS indicator H_2_DFFA was monitored for 495/520 nm (Ex/Em) 30 min in WT and Δ*pcrP* strains grown in BMM8 at 0 and 5 mM Ca^2+^ for 12 h. The change in fluorescence in 30 min was calculated for each replicate and was normalized by cell density. Light blue; 0 mM Ca^2+^, dark blue; 5 mM Ca^2+^. Two-way ANOVA; **** p-value ≤0.0001, * p-value ≤0.05

To gain structural insight into the interactions of PcrP, Acp3, and KatA, we applied computer modeling. This analysis not only supported the previously reported interaction between Acp3 and KatA (122) (Fig. S9B), but also predicted that the Acp3 binding site for PcrP overlaps with the binding site for KatA (Fig. S9C). This suggests that simultaneous binding of all three proteins would result in structural conflicts, making such an interaction unfavorable. While the precise role of PcrP in alleviating oxidative stress and its contribution to PMB resistance in *Pa* requires further experimental investigation, our findings support a model that PcrP bind and sequesters Acp3 from its interactions with KatA, thereby making the latter available for its enzymatic activity alleviating oxidative stress.

## DISCUSSION

Understanding microbial adaptations to the host environment is important for developing the means to control devastating bacterial infections, such as those caused by *Pa,* particularly in the lungs of patients with CF (124). Our previous work has shown that elevated Ca^2+^ levels, commonly encountered during CF disease (22,125,126), lead to significant proteomic and transcriptomic responses in *Pa* (19,21), resulting in increased virulence and tobramycin resistance (20,21,53). Here, we explored Ca^2+^ induction of *Pa* resistance to the “last resort” antibiotic PMB and identified three new genes contributing to Ca^2+^-dependent resistance to PMB. As summarized in figure 7, one of the genes, *PA2803* is transcriptionally regulated by Ca^2+^ and P_i_ starvation and therefore, was designated *pcrP*. We showed that PcrP lacks its annotated catalytic activity as phosphonatase and instead, interacts with different protein partners. Our data supported by computer modeling indicate that PcrP interactions with Acp3 may release catalase, KatA, and contribute to Ca^2+^-dependent protection from oxidative stress and thus PMB resistance. In addition, we show that PcrP is involved in *Pa’*s adaptations to P_i_ limiting conditions and contributes to the regulation of pyoverdine production. Since elevated Ca^2+^ and low P_i_ commonly coexist in the host during infections and control *Pa* virulence (11,26,31), PcrP may enable the integration of the two signals and provide the mechanistic basis for *Pa* adaptation to these conditions in the host.

**Figure 7.**
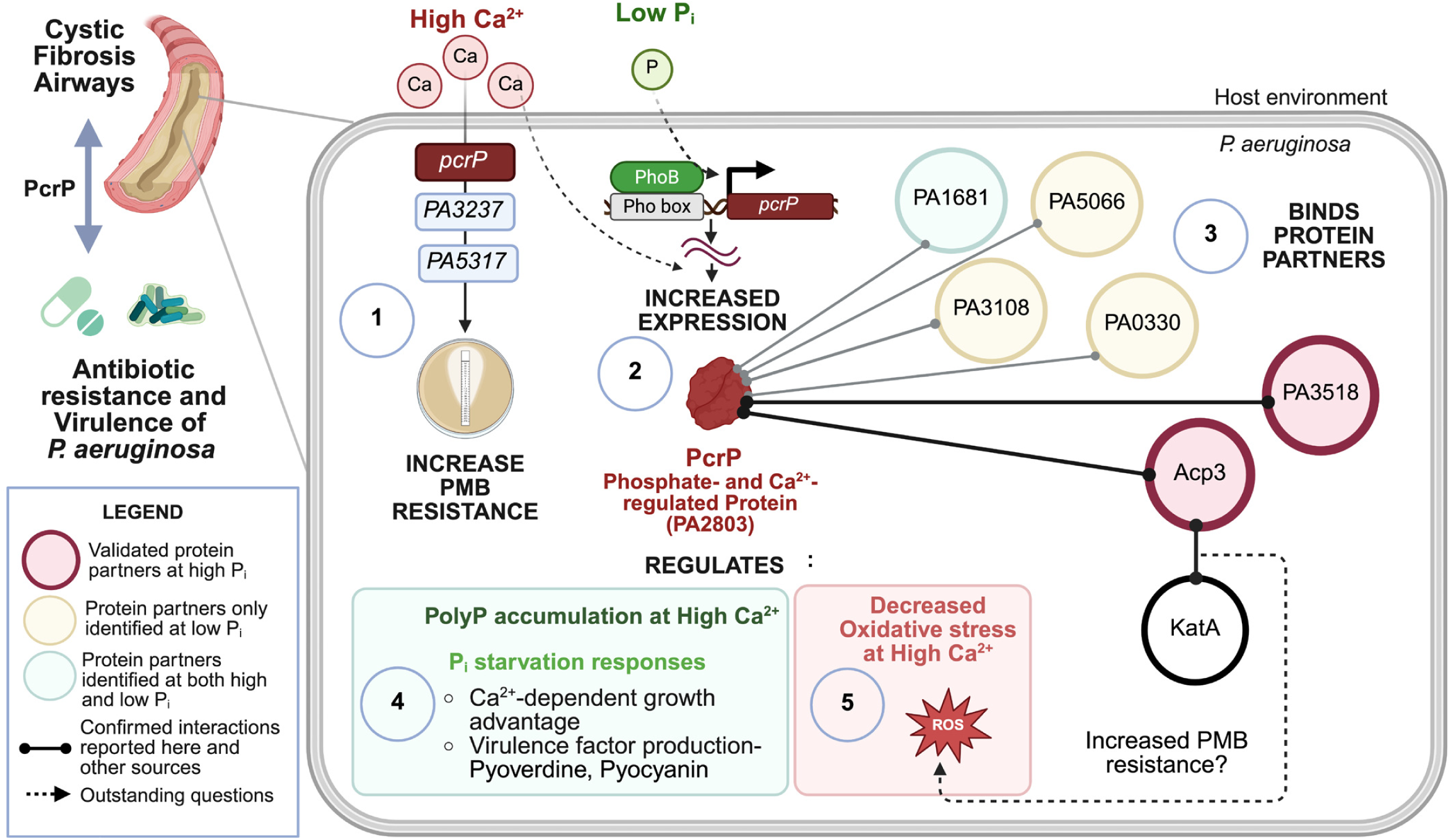
Summary of findings and proposed model illustrating the role of PcrP in Ca²⁺-regulated PMB resistance and P_i_ starvation responses in *Pa*. (**1**) *Pa* exhibits increased resistance to PMB under elevated Ca²⁺ through novel Ca²⁺-dependent mechanisms involving *PA2803* (herein named PcrP), *PA3237*, and P*A5317*. (**2**) Expression of *pcrP* is induced by elevated Ca²⁺ and P_i_ starvation, two host-associated environmental cues encountered during infection (24, 31, 114). (**3**) Catalytic assays and sequence analysis confirmed that PcrP lacks enzymatic activity and instead mediates protein-protein interactions, a feature likely conserved within the PA2803 subfamily of HADSF proteins. Several PcrP binding partners were identified, including proteins that bind specifically under high Ca²⁺ or P_i_-starved conditions, supporting PcrP role in *Pa* environmental adaptations. (**4**) PcrP contributes to the Ca²⁺-dependent *Pa* growth advantage under P_i_-limited conditions. PcrP also regulates polyP accumulation and modulates virulence factor production during P_i_ starvation. (**5**) PcrP interacts with Acp3, an inhibitor of KatA (122), which places PcrP within the regulatory network of oxidative stress in *Pa*. PcrP contributes to the Ca²⁺-dependent reduction in ROS production. Given the key role of oxidative stress in PMB’s bactericidal activity, PcrP-mediated mitigation of oxidative stress may contribute to the observed Ca²⁺-enhanced PMB resistance.

Several studies have shown that Ca^2+^ enhances the PMB resistance in *Pa* (89–91,127). However, the regulatory mechanisms of this resistance had not been identified. Consistent with the earlier reports (89,127), here we observed a 12-fold increase in PMB resistance in *Pa* strain PAO1 grown at 5 mM Ca^2+^ in comparison to that at no added Ca^2+^. As a first step in determining the responsible mechanisms, we examined the involvement of known PMB resistance mechanisms. Among several known regulators of PMB resistance in *Pa*, the TCSs PhoPQ, PmrAB, and ParRS have often been associated with elevated resistance observed in clinical strains (92–94). However, the mutations in these TCSs as well as in the Ca^2+^-responsive TCS, CarSR, and the efflux pump MexAB-OprM did not alter the Ca^2+^-induced PMB resistance in *Pa*. In addition, our transcriptional profiling showed that none of the 37 genes comprising the so far established PMB resistome in *Pa* (35,36,38,95,97,128) were positively regulated by Ca^2+^. In fact, 17 of them were downregulated (fold change ≤0.5) in the presence of 5 mM Ca^2+^. Notably, these downregulated genes included the TCSs and the *arn* operon, responsible for the covalent addition of L-Ar4N to lipid A of LPS, a key determinant of PMB resistance (128). These observations indicated that Ca^2+^-induced PMB resistance in *Pa* involves novel pathways. To identify genes involved in Ca^2+^-dependent PMB resistance, we used NTG mediated random mutagenesis followed by gene complementation and PMB susceptibility testing. This approach only allowed us to evaluate the role of non-essential genes in PMB resistance. We identified three genes: PA2803, PA3237 and PA5317, whose mutations resulted in impaired PMB resistance in *Pa* at elevated Ca^2+^. Considering PA2803 transcriptional regulation by Ca^2+^, this study focused on understanding the role of this gene, designated *pcrP,* encoding a putative phosphonatase belonging to HAD superfamily of proteins.

*Pa* responses to low availability of P_i_ are commonly detected during lung infections in CF patients, indicating that *Pa* encounters P_i_-starvation during CF lung colonization (31,129–131). In addition to Ca^2+^, the transcription of *pcrP* increases in response to low P_i_ under the control of PhoB, the global transcriptional regulator of P_i_ starvation adaptations in *Pa*, with a corresponding increase in PcrP abundance (31,113,132). Notably, *pcrP* transcription is also upregulated (log_2_ fold-change of 7.95) during biofilm formation (133) and in response to tobramycin treatment (134), indicating its broader role in the pathogen’s adaptation to infection-related conditions. The co-regulation of *pcrP* by multiple infection-related factors highlights its importance in *Pa’*s ability to thrive in host environments, such as CF lungs, where these conditions co-exist. Supporting this, metatranscriptomic analyses of CF sputum samples have shown *pcrP* to be highly expressed during infections (SRX5145606 and SRR6833349 with FPKM (Fragments Per Kilobase of transcript per Million mapped reads) values 12.0 and 4.18, respectively) (31,131,135). These findings highlight PcrP as a promising target for developing anti-virulent strategies aimed at modulating *Pa* virulence in CF and other disease settings.

According to our sequence analyses and *in-vitro* enzymatic assays, PcrP has neither phosphonatase nor phosphatase enzymatic activity. Instead, we identified an alternative protein-binding function for this protein, thus establishing an experimental precedent for a novel non-catalytic function in the PA2803 subfamily of HADSF proteins. Considering the phylogenetic conservation among the PA2803 subfamily members, lacking catalytic residues within their truncated cap domain (49), the protein-binding function is likely shared by all of them. The interactions of PcrP with PA3518 and Acp3 were validated by BTH and supported by computer simulations. The latter revealed that all three proteins can form stable homo- and heterodimeric complexes individually and simultaneously, though according to the energy analyses, the interactions of PcrP with Acp3 were more stable than with PA3518. While the MD simulations consolidate with our experimental observations, it is important to note they may not fully reflect the complexity of protein interactions in cells and therefore, the strength of these interactions may alter in the microenvironment within cells.

Although acyl carrier proteins (Acps) are traditionally known for their role in fatty acid synthesis (122,136), some bacterial Acps possess functions extending beyond this canonical role (122). For example, in *E. coli,* proteins such as IscS, MukB, and YbgC have protein-binding functions unrelated to fatty acid synthesis (122,137,138). Acp3 has been reported to interact with the major catalase KatA and repress its activity in *Pa* (122,139), though the physiological content and outcomes of this interaction remain unclear. Here we began exploring the role of PcrP-Acp3 interactions in Ca^2+^-dependent oxidative stress responses in *Pa*. Our results suggest that elevated Ca^2+^ induces a protective response in Pa against self-produced ROS, which involves PcrP. Such protective response is supported by our RNA-seq analysis showing that several oxidative stress genes, such as *sodM* (140) *ohr* (141), and *ahpC* (142), are upregulated at elevated Ca^2+^ (log_2_ fold-change values of 3.36, 3.18, 1.01, respectively). Furthermore, computer simulations showed that simultaneous binding of both PcrP and KatA to Acp3 appears unfavorable due to the overlap in the predicted binding regions, indicating that once PcrP binds to Acp3, KatA becomes available for its catalytic function. Since PMB increases ROS production and causes oxidative damage (143,144), this PcrP-dependent availability of KatA may play role in protecting *Pa* against the antibiotic at elevated Ca^2+^. In agreement, KatA has been shown to play a role in PMB-induced oxidative stress contributing to the intrinsic resistance observed in *Pa* (145). Further in support, the expression of *acp3* has been shown to increase in the presence of polymyxin-E (GEO profile 61228037), also supporting its importance for PMB resistance. Further studies are needed to characterize the molecular details of PcrP-Acp3 interactions and their role in regulating KatA catalase activity and Ca^2+^-enhanced PMB resistance in *Pa*.

A second protein partner of PcrP, PA3518, is encoded within an operonic gene cluster (PA3515-PA3519) regulated by a transcriptional regulator CueR, controlling Cu resistance in *Pa* (146,147). In addition, Cu stress responses in *Pa* involve CopRS, a second responder, activated upon prolonged Cu accumulation (148–150). According to (146), *pcrP* is regulated by CueR upon a short exposure to 0.5 mM Cu, whereas *PA3518* is regulated by CopR during prolonged Cu treatment (146). Moreover, the GEO profile (GDS2377) reports that both *pcrP* and *PA3518* are significantly upregulated in response to Cu shock (exposure to 10 mM for 45 min). This differential regulation of *pcrP* and *PA3518* by Cu stress suggests that their interaction may have a role in *Pa* protection against Cu toxicity. It is also noteworthy that PA3518 contains a heme oxygenase-like domain (InterPro entry: IPR016084), also found in such proteins as coenzyme PQQ (pyrrolo-quinoline-quinone) biosynthesis protein C (PqqC), an enzyme involved in producing the redox cofactor PQQ (151–154) and shown to enhance survival during oxidative stress in *Deinococcus radiodurans* (155). Cu stress is closely associated with oxidative stress, stemming from Cu propensity to produce ROS *via* auto-oxidation or Fenton-like reactions (149,156). This raises an interesting possibility that PA3518 may also be involved in oxidative stress responses in *Pa*. Furthermore, the promoter region of the operonic cluster of PA3518 harbors a Pho box, to which the regulator PhoB binds to and drives P_i_-starvation related responses (114), which suggests an additional role of PA3518 in P_i_ starvation responses.

Since the expression of PcrP is controlled by the availability of P_i_ (24), we predicted that the protein plays role in *Pa* responses to low P_i_. In fact, several HADSF proteins have previously been associated with P_i_ starvation responses in other organisms (157) owing to their wide range of enzymatic activities associated with phosphoryl transfer (158). Our findings indicate that elevated Ca^2+^ levels found in CF environment enhance the ability of *Pa* to survive P_i_ starvation and that PcrP is required for the full scale of this adaptation. To better understand the role of PcrP in P_i_ starvation responses, we followed the changes in P_i_ uptake, storage, and recycling in the deletion mutant of *pcrP*. The results indicate that among these, only production of poly P benefits from PcrP. Bacteria synthesize poly P complexes under a variety of conditions, especially during nutritional stress (116,159,160). These P_i_ reserves are broken down to replenish the cellular demand for P_i_ and allow bacteria to survive during starvation. It is possible that the Ca^2+^-dependent contribution of PcrP to poly P production provides the observed modest growth benefit during P_i_ starvation. Further, our results indicate Ca^2+^-dependent changes in the abundance of amino-lipids, such as OL, in *Pa*, biosynthesis of which has been correlated with PMB resistance (31). Further studies will follow the mechanistic link between this role of PcrP and the observed phenotypic differences in the mutant during P_i_ starvation.

In summary (Fig. 7), the identification of PcrP for its role in Ca^2+^-enhanced PMB resistance led to discovering its novel function as a networking protein, the function that is likely shared by the entire PA2803 subfamily of proteins. So far, the emerging functional network of PcrP links *Pa* responses to P_i_ starvation, ROS, Cu toxicity, and Ca^2+^-dependent PMB resistance and suggests that through protein-protein interactions, PcrP acts as a molecular hub integrating multiple environmental cues and mediating Ca^2+^-guided adaptation of *Pa,* ultimately enabling its survival during infections.

## ACKNOWLEDGEMENTS

**MP** acknowledges funding from NIH grants 1R15GM124670-0 and P20GM103648. **MJF** acknowledges funding from NIH grant NIH NIAID AI154171. **PKA** acknowledges funding from NIH/NIGMS (R01 grant GM148886) and computing time from Oklahoma State University’s High-Performance Computing Center funded by NSF awards (OAC-1531128 and OAC-2216084). We are grateful to Kerry Williamson from Montana State University for her help with MIC assessments of the mutant library. Protein MS/MS analysis was performed at the Oklahoma State University Proteomics Facility. We also acknowledge Brian Cougar (Oklahoma State University) for assisting with RNA-seq analysis and Dr. Tim Tolker-Nielsen (Copenhagen University, Denmark) for providing us *Pa* Δ*mexAB-oprM*.

